# A molecular logic of sensory coding revealed by optical tagging of physiologically-defined neuronal types

**DOI:** 10.1101/692079

**Authors:** Donghoon Lee, Maiko Kume, Timothy E Holy

## Abstract

Neural circuit analysis relies on having molecular markers for specific cell types. However, for a cell type identified only by its circuit function, the process of identifying markers remains laborious. Here, we report physiological optical tagging sequencing (PhOTseq), a technique for tagging and expression-profiling cells based on their functional properties. We demonstrate that PhOTseq is capable of selecting rare cell types and enriching them by nearly one hundred-fold. We applied PhOTseq to the challenge of mapping receptor-ligand pairings among vomeronasal pheromone-sensing neurons in mice. Together with *in vivo* ectopic expression of vomeronasal chemoreceptors, PhOTseq identified the complete combinatorial receptor code for a specific set of ligands, and revealed that the primary sequence of a chemoreceptor was an unexpectedly strong predictor of functional similarity.

---

Molecular markers have been a powerful tool for labeling and analyzing neuronal cell types. However, in many cases a single marker is insufficient to define a unique cell type, and may label a few, or a few hundred, physiologically-distinguishable cell types (*1*–*3*). In such cases, one wishes to select specific physiological populations and discover their molecular identities. However, tools for proceeding from function to molecular markers are not fully mature. For instance, activity markers such as *c-fos* allow identification of active neurons, but are difficult to leverage effectively in cases where single physiological types can only be defined via an intersectional approach involving multiple stimuli, experiences, and/or behaviors (*4*). Techniques like patch-seq enable expression-profiling of single recorded neurons (*5*–*8*); however, this approach is laborious, and faces daunting obstacles if one wants to collect dozens of examples of a particular rare cell type for deep molecular profiling of low-abundance transcripts.

To overcome these challenges, we reasoned that the key bottlenecks of this pipeline could be replaced by an all-optical approach. One prototype is CaMPARI (*9*), which uses the simultaneous presence of calcium (whose influx is triggered by neuronal activity) and excitation by short-wavelength light under the control of the experimenter. While CaMPARI allows one to control labeling of excited neurons temporally, as with *c-fos* any active neuron will be labeled, so that this may comprise dozens or more cell types. There is considerable need for tools that provide the specificity of patch-seq with the convenience and high-throughput of optical methods.

Here we report a systematic and comprehensive approach called physiological optical tagging sequencing (PhOTseq). PhOTseq allows specific cell types to be defined intersectionally via their physiological activity under diverse conditions. We exploit two coexpressed fluorescent proteins, GCaMP and photoactivatable mCherry (PAmCherry) (*10*–*12*): GCaMP permits recording neuronal activity by large-scale calcium imaging, and PAmCherry enables long-term labeling of selected neurons by photoactivation (PA). Using these reporters, one performs calcium imaging using a diverse set of stimuli or experiences, and then online analysis to select a specific collection of cells for tagging by photoactivation. Finally, the tagged neurons are harvested and profiled for their mRNA expression (**Fig. 1A**). To this basic PhOTseq workflow, we also added efficient methods for *post hoc* validation.

**Fig. 1.**
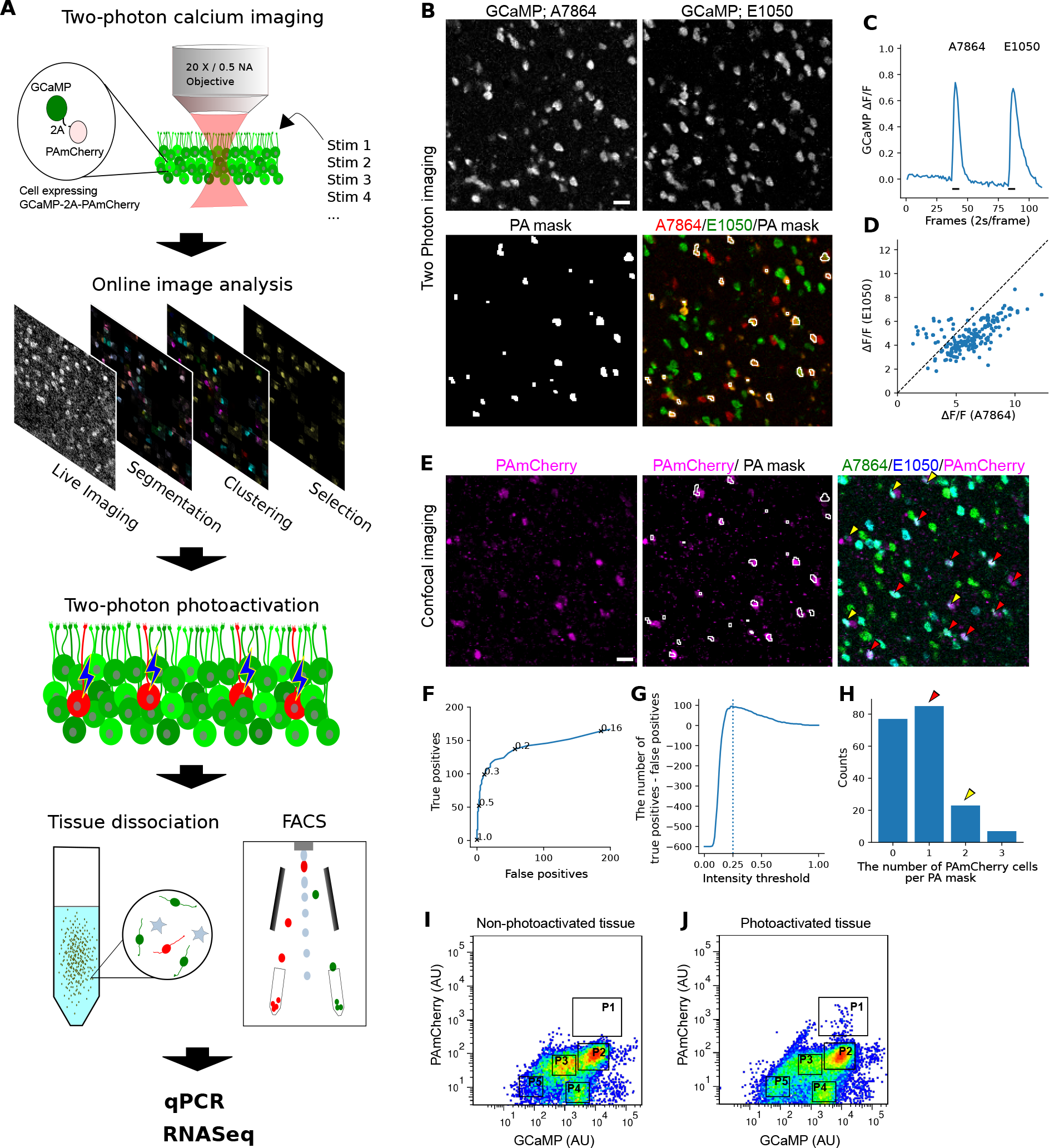
Two-photon photoactivation can tag neurons chosen by activity pattern. (A) The work flow of Physiological Optical tagging sequencing (PhOTseq). (B–D) Two-photon calcium imaging of the VNE explanted from a GCaMP5g-p2a-PAmCherry transgenic mouse. (B) Rep-resentative images of calcium chemosensory response evoked by either 10 *μ*M A7864 (top left) or 10 *μ*M E1050 (top right). For the cells responsive to both stimuli, photoactivation masks were drawn (bottom). Scale bar: 20 *μ*m. (C) The average GCaMP intensity was obtained from masked cells. Black bars below the trace signify delivery time-course of stimuli shown at the top. (D) Each dot indicates each masked cells’ response (Δ*F/F*) to 10 *μ*M A7864 and 10 *μ*M E1050. (E–H) After two-photon calcium imaging and photoactivation, PAmCherry signal and calcium responses were recorded using confocal microscopy. (E) The confocal images show PAmCherry signals (left), PAmCherry signals with two-photon PA mask (middle), and PAmCherry signals with calcium responses to either 10 *μ*M A7864 (cyan) or 10 *μ*M E1050 (green) (right). In the merged image (right), red and yellow arrow heads are the examples of PA mask regions associated with single and double PAmCherry-positive neurons, respectively. Scale bar: 20 *μ*m. (F) A receiver-operator characteric (ROC) analysis showed the number of true and false PA-positive regions at different PAmCherry intensity. The numbers on the ROC curve indicate PAmCherry detection threshold. PAmCherry intensity was normalized by the maximum PAmCherry intensity. (G) Differences between the number of true PA-positives and that of false PA-positives as a function of PAmCherry intensity threshold. PAmCherry intensity was normalized by the maximum PAmCherry intensity. The dashed line indicates 25% of the maximum PAmCherry intensity. (H) The histogram shows the number of PAmCherry-positive cells associated with each PA mask. Red and yellow arrow heads indicate PA masks associated with one and two PAmCherry positive cells, respectively; the example cells are shown in (D, red and yellow arrow heads). (I and J) Non-PA’ed (I) or PA’ed (J) tissues were dissociated and subjected to FACS analysis. Clusters were marked as P1, P2, P3, P4, and P5. (I) Few particles were detected in P1 (0.013%), while P2 (used for control cells) accounted for 33.9% of the total population. (J) VSNs responding to both A7864 and E1050 were selectively PA’ed and analyzed by FACS. A small number of particles was observed in P1 (0.218%). P2 contained 37.3% of the total population.

In developing PhOTseq, we created several plasmids, viruses, and a mouse line all expressing variants of GCaMP-2A-PAmCherry. The self-cleaving 2A peptide (*13*) allows both proteins to be expressed from a single mRNA; the tight coupling facilitates ratiometric analysis to compensate for variability in expression. Initial *in vitro* and *in vivo* experiments demonstrated the feasibility of performing both calcium imaging and photo-tagging using this reporter (**Fig. S1 and Fig. S2**). To test the performance of the complete pipeline—include near-realtime analysis of large population calcium imaging data and its use to select cells for photo-labeling (**Fig. 1A**)— we turned to mouse vomeronasal sensory neurons (VSNs), whose primary function is pheromone sensing and social communication. VSNs exhibit dense cell packing (*14*, *15*) (thus challenging photo-tagging accuracy), extreme functional heterogeneity (up to ~350 distinct cell types based on receptor gene expression), and an unambiguous relationship between molecular identity and physiological function (through expression of the receptor gene (*16*–*19*)). We created a transgenic tetO-GCaMP5g-2A-PAmCherry mouse and drove expression in all VSNs via OMP-IRES-tTA (**Fig. S3A**). We observed calcium responses from the VSNs, and two-photon photoactivation resulted in photo-tagging at single-cell resolution (**Fig. S3 and Fig. S4**).

To initially optimize and characterize high-throughput photoactivation of neurons chosen by their activity pattern, we evoked combinatorial VSN responses using two ligands, 5-androsten3*β*, 17*β*-diol disulphate (here called A7864) and 1, 3, 5(10)-estratrien-3, 17*β*-diol disulfate (E1050). A custom online algorithm automatically segmented responsive neurons in the data; out of 430 neurons responding to at least one ligand, 192 neurons responded to both ligands. Voxels corresponding to these dually-responsive neurons were chosen to create a “mask” so that only these voxels were illuminated during photoactivation (**Fig. 1, B-D**). After the ~7-minute photoactivation step, photoactivated neurons were clearly distinguishable from the background (**Fig. 1E**): As we increased the PAmCherry detection threshold, false PA-positives decreased rapidly while true PA-positives decreased slowly (**Fig. 1F**). At a threshold of 25% of the maximum PAmCherry signal, the number of true PA-positives exceeded the false PA-positives by the greatest margin, resulting in 115 true PA-positives and 20 false positives out of the 192 PA mask regions (**Fig. 1G**). We also analyzed the single-cell targeting precision within each PA mask region. Among true PA-positive regions, 85 (74%) were associated with single, 23 (20%) with two, and 7 (6%) with three PAmCherry^+^ cells (**Fig. 1H**). Therefore, an average of 1.32 cells per PA region were photoactivated. Given that such neurons comprised only 3–5% of the original population (see **Fig. 2B**), this demonstrates that PhOTseq is capable of greatly enriching rare types.

**Fig. 2.**
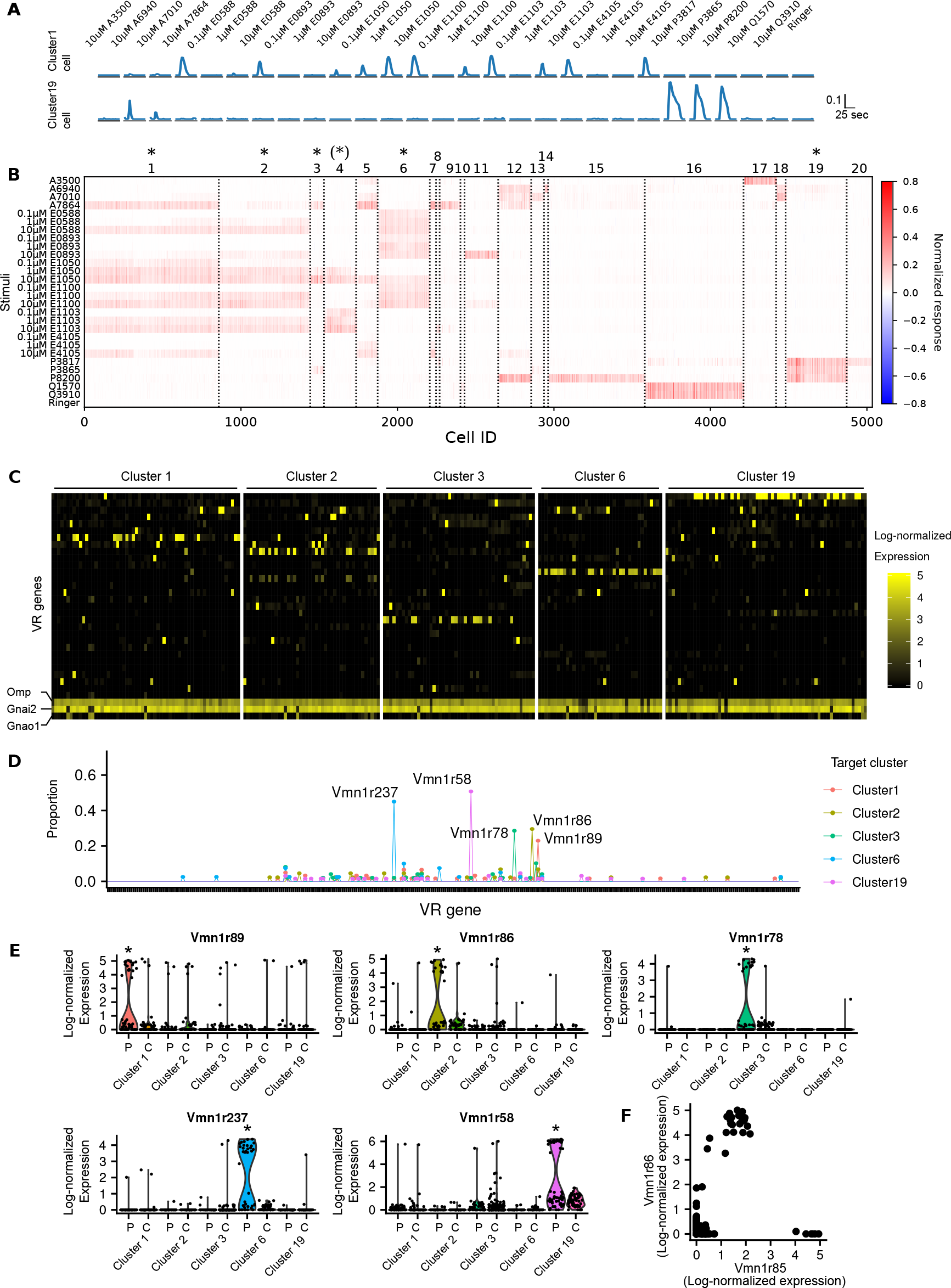
Physiologically similar cell types expressed distinct VR genes. (A) Example calcium traces from a cluster1 or a cluster19 cell. Amplitude was normalized by the maximum amplitude of all cells recorded. Ligands are listed at the top. (B) A heatmap showing neuronal responses to 15 different sulfated steroids, among which sulfated estrogens (six E compounds) included multiple concentrations (0.1 *μ*M, 1 *μ*M, and 10 *μ*M). If not written, the ligand concentration was 10 *μ*M. Cells are on columns, stimuli on rows. The color bar indicates normalized response (see Methods). Cluster identities are reported at the top. “*” marks PhOTseq target cell types. “(*)” marks the functional type whose receptor identity was discovered during analysis by ectopic expression in **Figure 3**. Responsive neurons were pooled from three imaging volumes (three VNEs). (C) A heatmap showing expression of the 30 most highly expressed VR genes when averaged across all sequenced cells; also shown are three marker genes (*Omp*, *Gnai2*, and *Gnao1*). Cells are on columns, genes on rows. PhOTseq-targeted functional types are shown at the top; in separate experiments, cells belonging to these functional clusters were specifically photoactivated and sequenced. The colorbar indicates log-normalized expression level. (D) The proportion of a VSN type in each group shown in (C). For each cell, the VSN type was defined as the maximally expressed VR gene. Each tick on the horizontal axis represents a different VSN type. Note that each functional type exhibits only one common VSN type. (E) Violin plots showing the expression of five VR genes across different experiment groups. “P” indicates photoactivated cells and “C” indicates non-photoactivated control cells. Asterisk(*) indicates *p*_adj_ < 0.01 (Wilcoxon rank-sum test; *p*_adj_: 2.7 *×* 10^−11^, 2.4 × 10^−10^, 1.1 × 10^−20^, 1.3 × 10^−40^, 2.7 × 10^−30^; average fold difference: 6.2, 23.8, 80.7, 75.7, 45.9 for Vmn1r89, Vmn1r86, Vmn1r78, Vmn1r237, Vmn1r58, respectively). (F) Single-cell expression of *Vmn1r85* and *Vmn1r86* is non-exclusive.

**Fig. 3.**
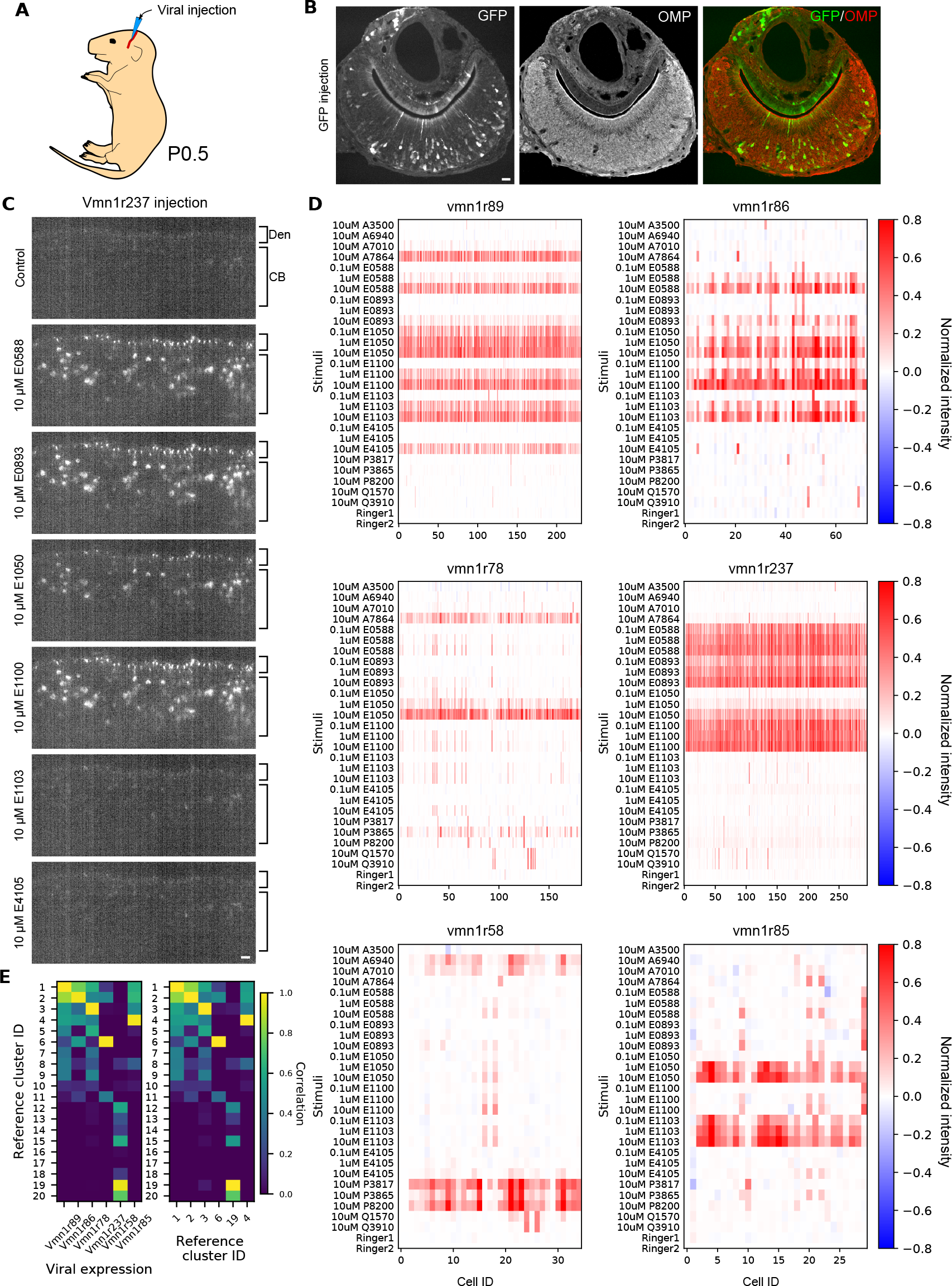
Ectopic expression allowed functional analysis of vomeronasal receptors. (A) A schematic of the temporal vein injection to a P0.5 mouse pup. (B) Coronal section of the vomeronasal organ after rAAV2/8-CAG-GFP transduction. The expression of OMP, which marks VSNs, was detected by immunohistochemistry. Scale bar: 20 *μ*m (C) An example optical section from light sheet calcium imaging after the ectopic expression of GCaMP-2A-Vmn1r237. The apical layer is occupied by dendritic tips (Den) with cell bodies (CB) below. Scale bar: 20 *μ*m (D) Calcium response after ectopic expression of GCaMP-2A-Vmn1r89, -Vmn1r86, -Vmn1r78, -Vmn1r237, -Vmn1r58, or -Vmn1r85. Ligands are identical to those in **Figure 2A and B**. For each ectopic expression calcium imaging experiment, a single mouse was used. (E) Pairwise correlation between the reference clustering shown in Figure 2B and the ectopic responses shown in (D) (left) or autocorrelogram of the reference clustering (right).

The photoactivated vomeronasal epithelia (VNEs) were subsequently subjected to dissociation and fluorescence-activated cell sorting (FACS; **Fig. 1, I and J**). As a control, we also sorted dissociated cells from non-photoactivated VNEs. In FACS analysis, 4–5 abundant clusters were observed, of which only one (P1) was almost entirely dependent upon photoactivation. This population represents the one selected later for sequencing. Altogether, these results showed that a physiological subset of neurons can be identified by calcium imaging, photoactivated, and collected at large scale.

Having demonstrated that PhOTseq can efficiently enrich specific functional types, we used it to begin unraveling the logic of chemosensation. Fewer than ten vomeronasal receptors (VRs) have ligands of known structure (*23*), and these are scattered widely across the gene family. We reasoned that a saturation analysis of VRs responsible for encoding a focused region of “chemical space” will provide new insights into the molecular logic of chemosensation. First, we examined functional types of VSNs broadly by light sheet calcium imaging while presenting 15 different sulfated steroids, of which six were also delivered over a range of concentrations, for a total of 27 different chemosensory stimuli. Approximately 15,000 neurons, ~10% of the total VSN population per hemisphere, were visualized in each imaging volume (*3*). In-depth offline analysis revealed that, from three imaging volumes, more than 5000 neurons responded to at least one of the ligands, and they fell naturally into 20 different response types (**Fig. 2, A and B**). For photo-tagging, we selected the most abundant cell type, cluster1, and three others showing similar chemoreceptive fields (cluster2, 3, and 6) (**Fig. S5, A and B**), among which cluster3 comprised fewer than 2% of the responsive neurons and presumably ~0.2% of the total VSN population (**Fig. 2B**). As a control group, we also chose one of the most dissimilar cell types, cluster19, which strongly responded to sulfated pregnanolones. Collectively, these classes provided a stringent test of our ability to resolve subtle differences and assess their correspondence to distinct molecular signatures.

Following two-photon calcium imaging and photoactivation (**Fig. S6 and Movie S1**), we collected photoactivated or non-photoactivated experimental control cells by FACS and performed single cell RNA sequencing (scRNAseq). We obtained a total of 622 qualified cells (**Fig. S7**), among which 61, 44, 49, 40, and 65 photoactivated cells were from photoactivation experiments aiming at functionally-defined cluster1, cluster2, cluster3, cluster6, and cluster19 cells, respectively (**Fig. 2C**).

Physiological responses of chemosensory neurons are thought to be largely determined by the VR genes they express (*20*). Therefore, our initial investigation was focused on the expression of VR genes. Among photoactivated cells, any given cluster exhibited substantial enrichment of at least one VR gene, with different VR genes enriched in different clusters (**Fig. 2C**). In contrast, among control cells VR gene expression was sporadic (i.e., not consistent across cells (**Fig. S8**)); because our targeting accuracy was less than 100% (**Fig. 1H**), we also expected some sporadic expression among cluster-selected photoactivated neurons (**Fig. 2C**). When the type of a VSN was defined by its maximally-expressed VR gene, only one abundant type was found among cluster-selected photoactivated neurons (**Fig. 2D**; cluster1: *Vmn1r89* type (25%); cluster2: *Vmn1r86* type (30%); cluster3: *Vmn1r78* type (27%); cluster6: *Vmn1r237* type (50%); cluster19: *Vmn1r58* type (58%)). These five VR genes were the only significantly enriched receptor gene in each PA’ed group compared to all the other cells (**Fig. 2E**). During these analyses, we updated the *Vmn1r237* gene model as the read coverage of *Vmn1r237* and cloning suggested transcript variants missing from existing gene models (**Fig. S9** and Data S1). We also unexpectedly observed coexpression of *Vmn1r85* in *Vmn1r86* neurons (**Fig. 2F and Fig. S10A**). Including data from our control and sporadic cells, coexpression was also observed for several other VR gene pairs, in each case consisting of a genomically-adjacent pair (**Fig. S10, B–E**). Taken together, we identified six putative VR genes mediating a focused set of responses.

To rigorously test whether PhOTseq identified true receptor-ligand pairs, we decided to perform a gain-of-function study. Due to abnormal localization of VR proteins (*21*), *in vitro* heterologous expression systems have had little success, particularly for the V1R family (*22*). We reasoned that a VSN cell’s endogenous machinery would allow functional expression of a VR gene, so, seeking a more efficient route than making a transgenic mouse (*6*), we explored a virus-mediated approach. Among viral vectors we tested, intravenous injection of rAAV2/8-CAG-GFP was able to induce expression of GFP in a subset of VSNs (**Fig. 3, A and B**). Next, we delivered rAAV2/8-CAG-GCaMP-2A-VR, where VR was one of the VR genes identified by PhOTseq. For each virus, we obtained multiple neurons showing statistically significant responses to at least one ligand (**Fig. 3C and Movie S2**). In each case, the dominant response pattern exquisitely matched the expected PhOTseq response pattern (**Fig. 3, D and E**). For example, the ectopic expression of *Vmn1r89*, which was identified as a cluster1 receptor, resulted in neurons responding to multiple sulfated estrogens and A7864, thus recapitulating the pattern of cluster1. Likewise, ectopic expression of *Vmn1r86*, *Vmn1r78*, *Vmn1r237*, and *Vmn1r58* induced response patterns strongly correlated with their expected functional clusters. In addition to these five receptors, we also tested *Vmn1r85*, as this gene was found to be coexpressed with *Vmn1r86* (**Fig. 2C**). Unexpectedly, the response pattern matched that of cluster4, which was another group similar to cluster1 but not chosen for analysis by PhOTseq (**Fig. 3, D and E**). Since we observed neurons expressing only *Vmn1r85* (**Fig. 2F**), our result suggested that the functional cluster4 cells were likely to express *Vmn1r85*. Taken together, these results validate the receptor-ligand pairings found by PhOTseq, and further demonstrate that a virus-mediated ectopic expression system can be used for functional analysis of VRs.

Only a few studies have examined the relationship between VR sequence and chemosensory function (*6, 23*), and never from the vantage point of complete knowledge of the genes that underlie a set of nearest neighbors in terms of function. To investigate this relationship, we first calculated pairwise distances among predicted V1R amino acid sequences, from which we created an unrooted phylogenetic tree (**Fig. 4A and B**). Among V1R genes expressed in functionally similar types, Vmn1r89, Vmn1r86, and Vmn1r85 were close to each other in the sense of putative evolutionary distance. On the other hand, two of the functionally similar cell types, Vmn1r78 and Vmn1r237, belonged to different branches, perhaps suggesting that they had diverged from the others at an earlier point in evolution. In this representation Vmn1r58, which was the chosen outlier in terms of function, was not notably more divergent from Vmn1r89, Vmn1r86, and Vmn1r85 than the more functionally-similar Vmn1r78 and Vmn1r237. Consequently this tree structure representation does not suggest a particularly close correspondence between function and primary sequence.

**Fig. 4.**
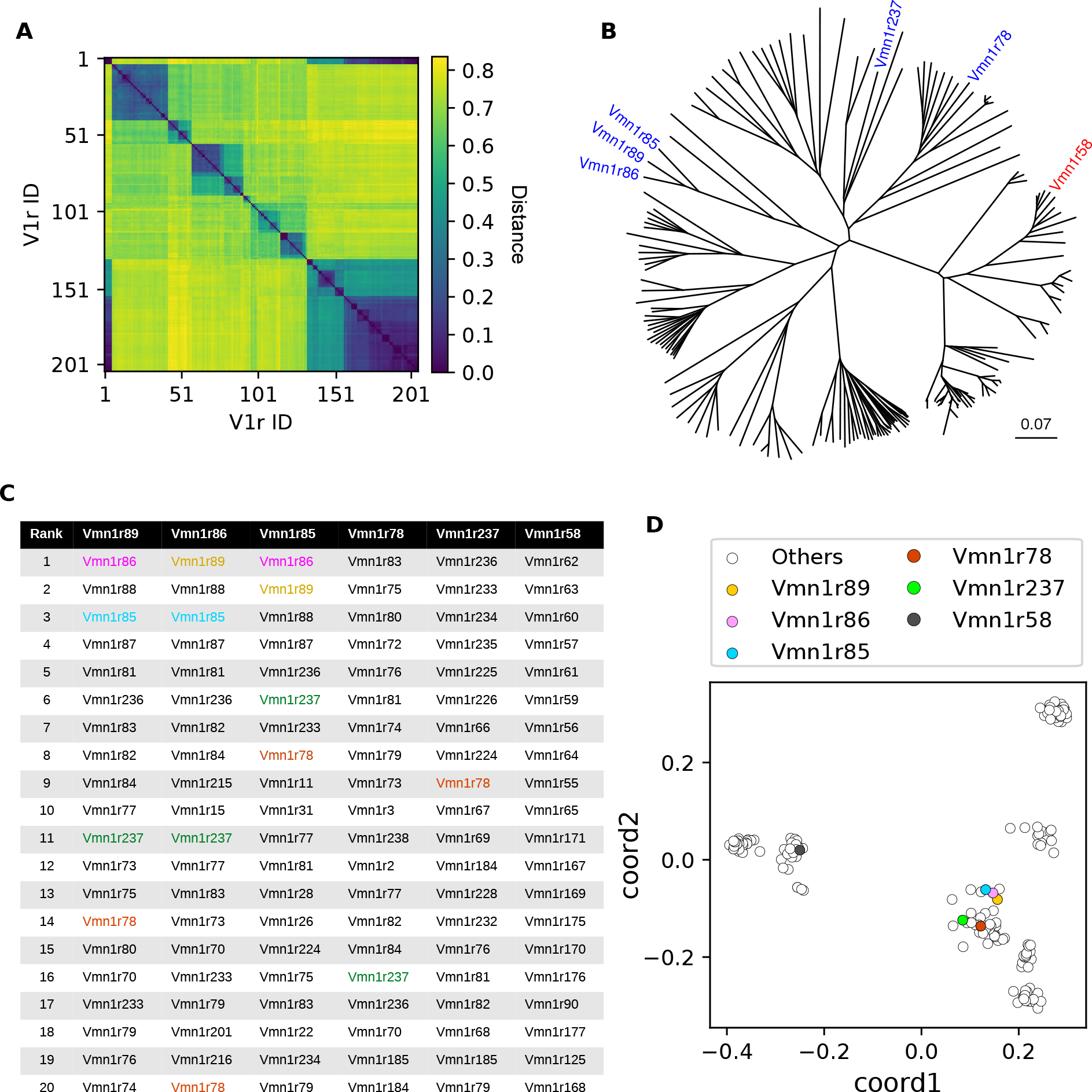
The similarities among sequences and among chemoreceptive fields are strongly correlated. (A) Pairwise distances among 205 V1R protein sequences (see Methods, (Sievers et al., 2011)). (B) An unrooted phylogenetic tree based on the distance matrix in (A). The five functionally similar (blue) and one distant (red) receptors studied here are marked. The scale bar indicates the number of amino acid substitutions per site. (C) For these deorphanized VRs, the top 20 nearest VRs, based on (A), were rank-ordered. The deorphanized VRs were colored if shown in the list (brown: Vmn1r89, pink: Vmn1r86, red: Vmn1r78, green: Vmn1r237, blue: Vmn1r85; Vmn1r58 was not shown in the rank table). (D) A 2-dimensional representation of the distance matrix in (A). Each dot represents a single VR gene.

However, when the distance matrix was examined carefully, we noticed that despite their apparent evolutionary distance, Vmn1r78 and Vmn1r237 were consistently among the most closely-related VR genes to Vmn1r89, Vmn1r86, and Vmn1r85 (**Fig. 4C**). Among 205 V1R genes examined, the median neighbor rank among these VRs was 11, implying that these genes are typically among the top few-percent closest matches to one another. Because distant leaves on the phylogenetic tree prove to be near-neighbors, this representation fails to convey the true proximity relationships among these sequences. To more comprehensively evaluate and visualize all pairwise distances, we therefore performed a classical multidimensional scaling analysis to project the high-dimensional sequence-based distance relationships into two dimensions (**Fig. 4D**). This resulted in a strikingly-different picture of the gene family, whose dominant feature was the presence of seven to nine apparent clusters. The functionally similar receptors, including Vmn1r78 and Vmn1r237, were near neighbors, whereas the functionally dissimilar receptor Vmn1r58 was positioned in a distant cluster. Using this more faithful representation of sequence-similarity, we conclude that relationships among the primary sequences of these VR receptors are strongly correlated with their degree of functional similarity.

Our results demonstrate the utility of PhOTseq, an all-optical solution to the problem of cell type selection and labeling. It provides an opportunity to study the relationships between genes and physiological function even in extremely rare cell types that can be defined only through extensive functional characterization. PhOTseq will be applicable in contexts in which individual neurons respond combinatorially to various conditions, including sensory stimulation, emotional status, and behavior. This work also presents *in vivo* ectopic expression, a new paradigm for functional VR gene expression. We anticipate that it will be applicable for characterizing VR and ligand pairings in detail. Lastly, our study provides biological insight on chemosensation by comprehensively mapping receptor-ligand pairings for chosen subsets of the vomeronasal sensory population, and suggests that saturation analyses can reveal sequence-function coupling unanticipated by single-ligand studies (*24*, *25*).

## Supporting information

Extended Database S1

## Acknowledgments

We thank Ron Yu for sharing the TetO vector and OMP-IRES-tTA mouse. We thank Paul H. Taghert for sharing the HEK cell line. We also thank Heide Schoknecht for mouse care and Dae Woo Kim for developing imaging software. Lastly, we thank Terra Barnes, Juntao Chen, Joseph Corbo, Xiaoyan Fu, Cody Greer, Alessandro Livi, Richard Roberts, and Manning Zhang for suggestions and comments. This work was supported by the Hope Center Viral Vectors Core and the Mouse Genetics Core at Washington University School of Medicine. This work was funded by NIH/NIDCD R01 DC005964 and DC010381.

## Supplementary materials

Materials and Methods

Figs. S1 to S10

References *(26-44)*

Movie S1

Movie S2

External Databases S1

## References

1. H. Markram, et al., Nature Reviews Neuroscience 5, 793 (2004).

2. M. Nassar, et al., Frontiers in Neural Circuits 9, 1 (2015).

3. P. S. Xu, D. Lee, T. E. Holy, Neuron 91, 878 (2016).

4. H. Okuno, Neuroscience Research 69, 175 (2011).

5. S. Qiu, et al., Frontiers in Genetics 3, 1 (2012).

6. S. Haga-Yamanaka, et al., eLife 2014, 1 (2014).

7. J. Fuzik, et al., Nature Biotechnology 34, 175 (2015).

8. C. R. Cadwell, et al., Nature Biotechnology 34, 199 (2015).

9. B. F. Fosque, et al., Science 347, 755 (2015).

10. J. Akerboom, et al., The Journal of Neuroscience 32, 13819 (2012).

11. T.-W. Chen, et al., Nature 499, 295 (2013).

12. F. V. Subach, et al., Nature Methods 6, 153 (2009).

13. M. D. Ryan, A. M. King, G. P. Thomas, Journal of General Virology 72, 2727 (1991).

14. K. Wilson, G. Raisman, Brain research 185, 103 (1980).

15. D. Keller, C. Erö, H. Markram, Frontiers in neuroanatomy 12 (2018).

16. T. Bozza, P. Feinstein, C. Zheng, P. Mombaerts, Journal of Neuroscience 22, 3033 (2002).

17. P. Feinstein, T. Bozza, I. Rodriguez, A. Vassalli, P. Mombaerts, Cell 117, 833 (2004).

18. E. A. Hallem, M. G. Ho, J. R. Carlson, Cell 117, 965 (2004).

19. X. Grosmaitre, A. Vassalli, P. Mombaerts, G. M. Shepherd, M. Ma, Proceedings of the National Academy of Sciences 103, 1970 (2006).

20. P. Mombaerts, Nature Reviews Neuroscience 5, 263 (2004).

21. J. Loconto, et al., Cell 112, 607 (2003).

22. B. Stein, M. T. Alonso, F. Zufall, T. Leinders-Zufall, P. Chamero, PLoS ONE 11, 1 (2016).

23. Y. Isogai, et al., Nature 478, 241 (2011).

24. B. von der Weid, et al., Nature Neuroscience 18, 1455 (2015).

25. Y. Jiang, et al., Nature Neuroscience 18, 1446 (2015).

26. P. S. Xu, T. E. Holy, Pheromone Signaling (Springer, 2013), pp. 201–210.

27. T. F. Holekamp, D. Turaga, T. E. Holy, Neuron 57, 661 (2008).

28. A. Kaur, S. Dey, L. Stowers, Pheromone Signaling (Springer, 2013), pp. 189–200.

29. H. A. Arnson, X. E. Fu, T. Holy, Journal of visualized experiments: JoVE 37 (2010).

30. T. Hashimshony, et al., Genome Biology 17, 1 (2016).

31. S. E. G. Lampe, B. K. Kaspar, K. D. Foust, Journal of visualized experiments: JoVE 93 (2014).

32. K. Kang, D. Lee, S. Hong, S.-G. Park, M.-R. Song, Journal of Biological Chemistry 288, 2580 (2013).

33. P. A. Gray, et al., Journal of Neuroscience 30, 14883 (2010).

34. T. E. Holy, Blockregistration package, https://github.com/HolyLab/ BlockRegistration (2019).

35. M. L. Comer, E. J. Delp, IEEE Transactions on image processing 9, 1731 (2000).

36. T. E. Holy, Cellsegmentation package, https://github.com/HolyLab/CellSegmentation (2019).

37. D. Turaga, T. EHoly, The Journal of Neuroscience 32, 1612 (2012).

38. T. E. Holy, Neighborhoodclustering package, https://github.com/HolyLab/NeighborhoodClustering.jl (2019).

39. A. Dobin, et al., Bioinformatics 29, 15 (2013).

40. S. Anders, P. T. Pyl, W. Huber, Bioinformatics 31, 166 (2015).

41. A. Butler, P. Hoffman, P. Smibert, E. Papalexi, R. Satija, Nature biotechnology 36, 411 (2018).

42. F. Hahne, R. Ivanek, Statistical Genomics (Springer, 2016), pp. 335–351.

43. F. Sievers, et al., Molecular systems biology 7, 539 (2011).

44. G. Yu, D. K. Smith, H. Zhu, Y. Guan, T. T.-Y. Lam, Methods in Ecology and Evolution 8, 28 (2017).

