## Supplementary Materials for "A molecular logic of sensory coding revealed by optical tagging of physiologically-defined neuronal types"

#### Materials and Methods

##### Animals

Experiments and animal care were performed in accordance with NIH guidelines and approved by the Washington University in St. Louis Institutional Animal Care and Use Committee (IACUC). Only healthy animals were used for this study. Mice were maintained on a 12hr:12hr light-dark cycle in a temperature-controlled facility. Mice were group-housed by sex from postnatal day 21–28 onward (P21–P28).

For ectopic expression, AAV injection was performed in P0–P1 male or female C57BL/6J mice. To obtain mouse pups, C57BL/6J mice were bred, and pregnant mice were visually identified and singly housed. After viral injection (see below), P0–P1 pups were returned to their mother until use or weaning at P21–P28.

For RNA in-situ hybridization and immunohistochemistry, 10–14-week old male C57/BL6J mice were used. Animals were singly housed one day before the experiment.

OMP-IRES-tTA mice (The Jackson Laboratory, 017754) were maintained by inbreeding OMP-IRES-tTA homozygous mice. TetO-GCaMP5g-P2A-PAmCherry mice (see below) were maintained by backcrossing with C57BL/6J and retaining pups containing the transgene. For calcium imaging, homozygous OMP-IRES-tTA mice and TetO-GCaMP5g-P2A-PAmCherry mice were bred to generate mice containing both *Omp-IRES-tTA* and *TetO-GCaMP5g-P2A-PAmCherry*. For light sheet calcium imaging, 10–16-week old male mice were used. For two-photon calcium imaging and PhOTseq, 10–16-week old male and female mice were used.

##### Generation of TetO-GCaMP5g-P2A-PAmCherry transgenic mouse

Construction of the TetO-GCaMP5g-P2A-PAmCherry transgenic line was outsourced to the Washington University in St. Louis mouse genetics core. A purified DNA construct (*TetO-GCaMP5g-P2A-PAmCherry*, see below) was injected into zygotes. Zygotes were obtained by mating B6CBAF1/J superovulated females with B6CBAF1/J males. The next morning, zygotes were collected from oviducts and microinjected in FHM embryo medium following standard procedures (1) with construct DNA at a concentration of 1.0 ng/$\mu$l, loaded into a single microinjection needle. For microinjection, the DNA was injected into the male pronucleus. Injections were performed with a Narishige micromanipulator and an Olympus inverted microscope using an air syringe as the injection device. Injected zygotes were transferred into pseudopregnant ICR female mice and live pups were born at day E19. Mice were handled according to institutional guidelines and housed in standard cages in a specific pathogen-free facility on a 14 h light/ 10 h dark cycle with ad libitum access to food and water. The existence of *TetO-GCaMP5g-P2A-PAmCherry* was confirmed by genotyping PCR (Table S1). A founder was backcrossed to C57BL/6J for more than five generations.

**Cell culture**

HEK293 cells were used for *in vitro* transfection. Cells were maintained in Dulbecco's Modified Eagle's Medium containing 10% fetal bovine serum in a humidified, 5% CO2 chamber at 37°C. Cells were subcultured when the confluency was approximately 80% in a 100 mm $\times$ 20 mm culture dish.

To make AAV (see below), the packaging cell line, HEK293, was maintained in DMEM supplemented with 5% fetal bovine serum (FBS), 100 units/ml penicillin, 100 $\mu$g/ml streptomycin in 37°C incubator with 5% CO2. The cells were plated at 30–40% confluence in CellSTACS (Corning, Tewksbury, MA) 24 h before transfection (70–80% confluence when transfection).

**DNA construction**

*GCaMP5g* was amplified by PCR from a plasmid purchased from Addgene (Plasmid #31788). The double-stranded *P2A* DNA was created by annealing sense and antisense *P2A* DNA strands (synthesized from IDTDNA; Table S1) containing EcoR1 and BamH1 target sequences on opposite 5' ends. 5.55 $\mu$M of each strand was mixed, heated up to 95°C, and slowly cooled to room temperature (RT). The 5'-ends of the resulting fragments were phosphorylated by T4 polynucleotide kinase (NEB, M0201S) in preparation for the ligation reaction. To create *GCaMP5g-P2A-PAmCherry*, the resulting *GCaMP5g* and *P2A* DNA fragments were ligated into a pPAmCherry1-N1 plasmid (Clontech, 632584). This plasmid was amplified using One Shot Top10 chemically competent E.Coli (Thermo Fisher Scientific, C404010). Depending on experimental purposes, *GCaMP5g* and *PAmCherry* were replaced with *GCaMP6f* and several VR genes, respectively, by GIBSON assembly (SGI-DNA, GA1100-10), and the resulting DNA constructs were amplified and subcloned into other vectors: pTRE2-GCaMP5g-P2A-PAmCherry was used for creating a transgenic mouse line. pAAV-CAG-GCaMP5g-P2A-PAmCherry and pAAV-CAG-GCaMP6g-P2A-PAmCherry were created for *in vitro* transfection and viral production. pAAV-CAG-GCaMP5g-P2A-VR and pAAV-CAG-GCaMP6f-P2A-VR, where VR is one of *Vmn1r89*, *Vmn1r86*, *Vmn1r85*, *Vmn1r78*, *Vmn1r237*, and *Vmn1r58*, were used for ectopic viral expression. The plasmids used for *in vitro* transfection and viral production were amplified by XL10-Gold competent E.Coli (Agilent Technologies, 200315)

The transgene construct was processed as follows: pTRE2-GCaMP5g-P2A-PAmCherry, which contains a tetracycline operator (TetO) promoter, was linearized by AatII and AseI. The DNA fragment containing *TetO-GCaMP5g-P2A-PAmCherry* was gel-purified by Qiaex II (Qiagen, 20021).

##### *In vitro* calcium imaging and photoactivation

*GCaMP5g-P2A-PAmCherry* was subcloned into an pAAV plasmid vector, which was kindly provided by the Washington University Hope Center Viral Vectors Core. The resulting plasmid was transfected into HEK293 cells using a commercial transfection reagent, Lipofectamin 2000 (ThermoFisher Scientific, 11668027). Two days after transfection, calcium imaging and photoactivation was performed by an epi-microscope (Leica microscope, DMI6000 B; Excelitas technologies light source, X-Cite exacte; HAMAMATSU CCD camera and controller, ORCA-R2/C10600). During calcium imaging, cells were continuously superfused with culture media, Dulbecco's modified Eagles medium (DMEM; ThermoFisher Scientific, 11995065). To transiently evoke a calcium response, a culture medium containing 10 $\mu$M ATP (Thermo Fisher Scientific, R0441) was delivered for 10 seconds. To increase intracellular calcium concentration, a culture medium containing 5 $\mu$M Ionomycin (Abcam, ab120116) was applied. Photoactivation was provided up to 5 minutes by a 405 nm laser at 1 mW output from the objective.

##### Recombinant AAV vector production

Recombinant AAV vector production was outsourced to the Washington University Hope Center Viral Vectors Core. 730 $\mu$g pAAV2/8 (for AAV8) or pAAV2/1 (for AAV1), 1180 $\mu$g pHelper, and 590 $\mu$g rAAV transfer plasmid containing the gene of interest were co-transfected into HEK293 cells, using the calcium phosphate precipitation procedure (Zolotukhin, et al 2002). The cells were incubated at 37°C for 3 days before harvesting. The cells were lysed by three freeze/thaw cycles. The cell lysate was treated with 25 U/ml of Benzonase at 37°C for 30 min followed by iodixanol gradient centrifugation. The iodixanol gradient fraction was further purified by column chromatography using a HiTrap Q column (GE

Healthcare). The eluate was concentrated with Vivaspin 20 100K concentrator (Sartorius Stedim, Bohemia, NY). Vector titer was determined by dot blot assay.

##### *In vivo* viral injection and photoactivation

*GCaMP6f-P2A-PAmCherry* was subcloned into an pAAV transfer plasmid, which was kindly provided by the Washington University Hope Center Viral Vectors Core. The resulting plasmid was sent to the viral core, and rAAV2/1-CAG-GCaMP5g-P2A-PAmCherry was produced. The virus was injected into the mouse somatosensory cortex: A C57BL/6J mouse was anesthetized with isoflurane (0.5–1.5%) while keeping its body temperature at 37°C. The scalp was clipped and cleaned using a surgical scrub. A cranial window was created over the cortex by a dental drill mounted on the stereotactic frame. A glass pipette with a tip diameter of 10–20 $\mu$m was slowly lowered into the brain, and 50–200 nl of virus was pressure-injected. After injection, the needle was slowly removed, and a coverglass placed over the hole. To head-fix the mouse during imaging, a headplate was attached to the skull by dental cement. After three to five weeks of recovery, the mouse underwent two-photon imaging and photoactivation. The mouse was head-fixed and placed on a cylindrical treadmill that allows the mouse to move while fixed in place. For *in vivo* imaging and photoactivation, we used a NA 1.0 objective (Olympus, XLUMPlanFLN20XW). *GCaMP6f*-expressing cells were visualized with a 910 nm two-photon excitation laser. Photoactivation was provided by a 810 nm Mai Tai DeepSee laser at 150 mW 1000–1500 "frames" over a cell body region, using 0.8 $\mu$s/pixel dwell time.

##### VSN ligands: sulfated steroids

Sulfated steroids used in our experiment were purchased from Steraloids: A3500: 5$\beta$-androstan-3$\alpha$-ol-11, 17-dione, sulphate, sodium salt; A6940: 4-androsten-17$\alpha$-ol-3-one sulphate sodium salt; A7010: 4-androsten-17$\beta$-ol-3-one sulphate, sodium salt; A7864: 5-androsten-3$\beta$, 17$\beta$-diol disulphate, disodium salt; E0588: 1, 3, 5(10), 7-estratetraen-3, 17$\beta$-diol 3-sulphate, sodium salt; E0893: 1, 3, 5(10)-estratrien-3, 17$\alpha$-diol 3-sulphate, sodium salt; E1050: 1, 3, 5(10)-estratrien-3, 17$\beta$-diol disulphate, disodium salt; E1100: 1, 3, 5(10)-estratrien-3, 17$\beta$-diol 3-sulphate, sodium salt; E1103: 1, 3, 5(10)-estratrien-3, 17$\beta$-diol 17-sulphate, sodium salt; E4105: 4-estren-17$\beta$-ol-3-one sulphate, sodium salt; P3817: 5$\alpha$-pregnan-3$\alpha$-ol-20-one sulphate sodium salt; P3865: 5$\alpha$-pregnan-3$\beta$-ol-20-one sulphate, sodium salt; P8200: 5$\beta$-pregnan-3$\beta$-ol-20-one sulphate, sodium salt; Q1570: 4-

pregnen-11$\beta$, 21-diol-3, 20-dione 21-sulphate, sodium salt; Q3910: 4-pregnen-11$\beta$, 17, 21-triol-3, 20-dione 21-sulphate, sodium salt

To make stock solutions, these sulfated steroids were dissolved in 50% methanol at 10 mM concentration, and they were stored in either 4°C or -20°C. To make the working concentration (0.1–10 $\mu$M), each stock was diluted in fresh carboxygenated Ringer's solution. Ligands were delivered to the VNE using an HPLC pump and an autosampler (Thermo scientific, Ultimate3000 pump and autosampler).

#### Calcium imaging of the vomeronasal sensory epithelium

Intact mouse VNEs were dissected out, flattened, and attached to a nitrocellulose membrane as previously described (*26*). Every step was done in ice-cold carboxygenated Ringer's solution (115 mM NaCl, 5 mM KCl, 2 mM $CaCl_2$, 2 mM $MgCl_2$, 25 mM $NaHCO_3$, 10 mM HEPES, and 10 mM freshly added D-(+)-glucose) Then, the epithelium-nitrocellulose membrane complex was mounted in an imaging chamber for light sheet calcium imaging or two-photon calcium imaging and photoactivation. To stabilize the tissue, it was incubated in ice-cold Ringer's solution for at least 30 min and then at imaging temperature for approximately two hours.

For light sheet calcium imaging, 620 $\mu$m $\times$ 640 $\mu$m $\times$ 300 $\mu$m imaging volumes were acquired at 0.5 Hz using a custom Objective-Coupled Planar Illumination (OCPI) microscope ( (*27*) retrofitted with a PCO edge5.5 sCMOS camera). Ringer's solution was superfused constantly at a rate of 540 $\mu$l/min at 34°C. To measure VSN responses, we applied 15 different sulfated steroids at 10 $\mu$M, among which six E compounds also included two other concentrations, 0.1 $\mu$M and 1 $\mu$M. In addition, two different aliquots of Ringer's solution were delivered as controls. Each stimulus was presented once for 5.5 seconds in each stimulus block; there were 4 stimulus blocks total, each presenting the ligands in different pseudo-randomized order interleaved with the other stimuli.

In the case of two-photon calcium imaging, 550 $\mu$m $\times$ 550 $\mu$m images were acquired at 0.5 Hz at room temperature. We used a 0.5 NA objective (Leica, 506147) with a pixel size of approximately 1.28 $\mu$m $\times$ 1.28 $\mu$m. Ringer's solution was consistently superfused at 540 $\mu$l/min. Among the fifteen different sulfated steroids used in the light sheet calcium imaging, five or six were chosen for the two-photon experiments. Each ligand was presented once for 4 seconds.

##### Two-photon photoactivation

After two-photon calcium imaging, responsive neurons were segmented by a custom segmentation algorithm written in the Julia programming language (see below). Then, a specific physiological cell type was selected under human control by iteratively thresholding GCaMP responses until a two-dimensional PCA scatterplot of selected ROI's responses appeared to be a single "blob." We obtained coordinates of the selected cells using a custom script written in Julia and created masks that were passed to the microscope software (Prairie view), so that the two-photon laser excites only pixels highlighted by the mask. Photoactivation was provided by a 810 nm Mai Tai DeepSee laser at 100–150 mW, scanning 400 "frames" with 0.8 $\mu$s/pixel dwell time for a total of approximately 5 minutes photoactivation time per field-of-view. An objective with 0.5 NA was used (Leica, 506147) and the pixel size was chosen to be 1.28 $\mu$m $\times$ 1.28 $\mu$m. The tissue depth at which photoactivation was applied ranged from 50–100 $\mu$m.

##### Manual photoactivation accuracy test

A VNE expressing *GCaMP5g-P2A-PAmCherry* in its VSNs was subjected to two-photon photoactivation. Cells were visualized by 910 nm excitation laser at 10–15 mW. Using the microscope software (Prarie view), photoactivation masks were manually drawn over the centers of a few arbitrarily-selected neurons (approximately 5 $\times$ 5 pixels). Then, photoactivation was provided by a 810 nm Mai Tai DeepSee laser at 100–150 mW scanned for 800 frames using 0.8 $\mu$s/pixel dwell time. An objective with 0.5 NA (Leica, 506147) was used and the pixel spacing was 0.64 $\mu$m. After photoactivation, 3d images were taken using a NA 1.0 objective with a pixel spacing of 0.57 $\mu$m and a $z$-step size of 1 $\mu$m. To reduce noise, images were averaged over 32 frames. PA target cells and cells nearby were identified by PAmCherry signal and the baseline GCaMP intensity. Then, they were manually segmented. Their positions and intensities were extracted using Fiji and analyzed in Julia.

##### Automated photoactivation accuracy test

A VNE expressing *GCaMP5g-P2A-PAmCherry* was subjected to two-photon calcium imaging. Two ligands, 10 $\mu$M A7864 and 10 $\mu$M E1050 were sequentially presented. After automatic cell segmentation (see below), neurons responsive to both ligands were chosen for photoactivation. Photoactivation was provided as described above for 400 scanned frames. After two-photon photoactivation, the tissue was moved to a confocal microscope where PAmCherry signals and GCaMP signals evoked by the two ligands were recorded. To find the photoactivated region, PAmCherry signals were used as markers, yet there noticeable differences between two-photon images and confocal images. Therefore, we registered two-photon images to confocal images by the following way: First, we manually-identified four cells that appeared both in two-photon and confocal calcium images. Second, the two-photon images were affine-transformed to match these cells. Lastly, the two-photon images were further registered by affine transformation to minimize the mean-square error between the two-photon images and the confocal images. The resulting transformation matrix was also applied to the two-photon PA mask image.

From a confocal PAmCherry image, we detected cells by the following way: The PAmCherry image was filtered by a Gaussian filter followed by a negative Lapacian of Gaussians filter using Julia (Images.jl package). Then, local maxima were detected in Fiji. Some local peaks were located in the middle of two or three adjacent cells. In such cases, we manually adjusted the position of local maxima to the center of one cell and added new marks to the centers of the other cells. We initially identified more than 1300 peaks, mostly based on the baseline PAmCherry intensity, and regarded these peaks as the centers of cells. The positions of these peaks were extracted using Fiji and further analyzed in Julia. The intensity of each cell was calculated by averaging pixel intensities around the peak. Then, we defined a true PA-positive region as all cells closely located to a PA mask ($< 15\mu$m). A false PA-positive region was defined as a cell distant from every PA mask. As a result, we initially obtained 189 true PA-positive out of 192 PA regions, 3 false PA-negative (192-189), and 789 false PA-positive regions. We changed the PAmCherry detection threshold to maximize the number of true PA-positives after subtracting the number of false PA-positives. At around 25% of the maximum PAmCherry signal, the number of true PA-positive (115) regions most outweighed false PA-positives (20), resulting in the sensitivity and specificity of 0.6 (115/192) and 0.97 (769/789), respectively. We reasoned that the 77 false PA-negative regions ($192 - 115$) were possibly due to different imaging conditions (e.g., out-of-plane localization), and FACS analysis might not be affected by such issues. In addition, PA masks at these false-negative locations tended to be smaller, suggesting that some of small PA masks did not efficiently induce photoactivation (data not shown).

#### Cell dissociation

We dissociated VSNs as described (*28*) with minor modifications: To activate papain, 1.1 ul of 1 M DL-Cys-HCl, 2.2 ul of 0.1 M EDTA, and 0.484 units of papain was dissolved in 200 ul of Hibernate A -Calcium solution (BrainBits, HACA), and incubated at 37°C for 30 minutes. During Papain activation, the basal lamina was removed from the VNE using a razor in ice cold Ringer solution (*29*). Then, VNEs were moved to a 1.5 ml Eppendorf tube and treated with 200 $\mu$l of the activated papain for 20 minutes at 37°C with gentle rotation. To this reaction, we added 200 $\mu$l of PBS (with calcium and magnesium; Gibco, 14040117)) containing 10 units of DNase. This reaction was further incubated for 5 minutes at 37°C with gentle rotation. At this point, we observed that cells were almost dissociated. Cells were further dissociated by pipetting 5–7 times: A Pasteur glass pipette was fire-polished to make an inner tip diameter of 150–200 $\mu$m and then coated with BSA (2.5 mg/ml; Jackson ImmunoResearch Laboratories, Inc., 001-000-161) by pipetting just before use to prevent cell adhesion. For washing, 1 ml of Hibernate A Low Fluorescence media (BrainBits, HALF) was added to make 1.4 ml. Cells were centrifuged for 3–4 minutes at 150 g, and half of the supernatant was replaced by new media. After centrifugation, half of the supernatant was removed to make $\sim$800 $\mu$l. Cells were re-suspended by gentle pipetting, filtered with a 40 $\mu$m cell strainer (Scienceware Flowmi, BAH136800040-50EA), and collected into a new tube.

#### Fluorescence-activated cell sorting (FACS)

We collected dissociated cells using FACS. Sorting gates were manually drawn by human judgment: Particles showing relatively lower SSC-A and FSC-A were discarded as they usually indicate cellular debris. We discarded outliers with high FSC-W and SSC-W as they possibly indicate doublets. Particles showing PAmCherry$^{high}$ and moderate GFP signal (P1 in **Fig. 1J**) were collected as photoactivated cells since this population did not appear in non-photoactivated tissues. Particles with PAmCherry$^{mid}$ and moderate GFP signal (P2 in **Fig. 1J**) were collected as experimental control cells. Relatively smaller number of particles showed considerably high GCaMP intensity. This is likely due to either cell damage or a spontaneous increase in intracellular calcium concentration during FACS. We did not collect these cells.

**Single cell RNAseq library prep**

We followed Cel-seq2 protocol (*30*) with minor modifications. Briefly, single cells were collected into a 96-well PCR plate containing 1.4 $\mu$l of lysis buffer in each well. The lysis buffer consisted of 0.096% Triton X-100, 0.08 U RNase inhibitor (Promega, N2615), 1 nmol dNTP and 0.2 pmol of reverse transcription primer. After collecting cells, plates were sealed and stored at -80°C until use. Subsequent steps were implemented as described in the original Cel-seq2 protocol. Seventeen sequencing libraries, altogether from 673 cells, were subjected to a 26bp $\times$ 98bp paired-end sequencing on two sequencing lanes of HiSeq2000.

**VR ectopic expression**

cDNA was extracted from the VNE using TRIzol (ThermoFisher Scientific, 15596026), and multiple VRs (*Vmn1r89*, *Vmn1r86*, *Vmn1r85*, *Vmn1r237*, *Vmn1r78*, and *Vmn1r58*) were amplified by PCR (Table S1). Each VR gene was ligated with *GCaMP* (either *GCaMP5g* or *GCaMP6f*) to make *GCaMP-P2A-VR* (see above). The resulting construct was subcloned into the pAAV transfer vector (see above). Then, rAAV2/8-CAG-GCaMP-P2A-VR was produced by the Washington University Hope Center Viral Vectors Core. A 50 ul aliquot of $10^{12}$–$10^{13}$ vg/ml virus was injected into the temporal vein of C57BL/6J P0.5 mouse (*31*). One to four weeks after viral injection, the VNEs were dissected out and prepared for light sheet calcium imaging.

***In vivo* sensory exposure and tissue sectioning**

For Fig. S10A, 10–14 wk old mice were singly housed for one day in a fresh cage. On the following day, the mouse was moved into another fresh cage. Sulfated steroid stock solution (10 mM, see above) was diluted in PBS to make 50 $\mu$M. Holding the mouse by hand, 10 $\mu$l of ligand solution was directly spotted dropwise onto the nostrils. With each mouse, this was repeated five times with the same stimulus, with a 5-minute gap between applications. Thirty minutes after the last treatment, the mouse was euthanized and the VNOs dissected out. The VNOs were fixed in 4% paraformaldehyde (PFA) overnight at 4°C. Subsequently, they were briefly rinsed with PBS, and incubated in 30% sucrose overnight. These were embedded in O.C.T. compound (Sakura Finetek, 4583) and stored at -80°C until cryosectioning. Cryosectioning was performed at -20°C. The thickness of each section was 12 $\mu$m. Sections were attached onto glass slides and stored at -20°C until use.

##### *In situ* hybridization and immunohistochemistry

Riboprobes were synthesized by *in vitro* transcription: The full-length coding sequence of VR genes (*Vmn1r85* : NM_145847.1; *Vmn1r86* : NM_001167536.1) were PCR amplified with the T7 promoter (GTAATACGACTCACTATAGG) attached at the 3'-end. The PCR product was purified (Qiagen, 28706) and used as DNA template (~500 ng) for *in vitro* transcription. In an *in vitro* transcription reaction (Promega, P2075), either DIG RNA labeling mix (Sigma-Aldrich, 11277073910) or Fluorescent RNA labeling mix (Sigma-Aldrich, 11685619910) was added. The transcription reaction was incubated at 37°C for 2 hours. Then, 2 units of DNase were added and incubation was further continued for 10 minutes at 37°C. To stop the transcription reaction, we added pH8.0 EDTA to a final concentration of 10 mM. Riboprobes were purified by LiCl precipitation, dissolved in $H_2O$, and stored in -80°C until use.

Fluorescent *in situ* hybridization was performed as described (Perkin elmer, NEL744001KT; (*32, 33*)) with modifications: Here, if not specified, the temperature was RT. Tissue sections were air-dried more than 15 minutes and fixed with 4% PFA for 20 minutes. Then, they were washed twice with PBS for 5 minutes each, and treated with 1% $H_2O_2$ in PBS for 1 hour. Next, they were rinsed with PBS twice for 5 minutes each, and incubated in RIPA buffer for 1 hour. After two 5-min PBS washings, they were fixed again in 4% PFA for 15 minutes. Following PBS washing (5 minutes, twice), they were acetylated with 1% acetic anhydride in 0.1 M triethanolamine-HCl, pH 8.0 for 10 minutes. After washing with PBS (3 minutes, 3 times), they were incubated with hybridization solution (50% formamide, 5X SSC, 0.3 mg/ml yeast RNA, 0.1 mg/ml heparin, 1X Denhardts, 0.05% Tween 20, 5 mM EDTA) for 1–4 hours at 65°C. Then, the hybridization solution was replaced with new hybridization solution containing ~2.5 $\mu$g/ml of riboprobes. The tissues were further incubated at 68°C overnight. On the following day, the slides were washed with pre-heated 5× SSC (10 minutes) and 0.2× SSC (1 hour, twice) at 70°C. Then, they were washed with 0.2× SSC (5 minutes) and with 0.1% PBST (PBS + 0.1% Triton X-100; 5 minutes) at RT. The tissues were treated with 1× blocking solution (10× blocking solution (Sigma-Aldrich, 11096176001) diluted in 0.1% PBST) for an hour at RT. The solution was replaced with new blocking solution containing anti-Fluorescein-POD antibody (1:2500; Sigma-Aldrich, 11426346910), anti-pS6 (240/244) antibody (1:500; Cell signaling technology, 2215S), and anti-pS6 (244/247) antibody (1:1000; ThermoFisher, 44923G), and incubated overnight at 4°C. On the third day, the slides were washed with 0.1% PBST (5 minutes, four times), and treated with Fluorescein-conjugated TSA (Perkin elmer, SAT701001EA) for 5–20 minutes. They were washed with 0.1% PBST (5 minutes, four times) and incubated in 3% $H_2O_2$ for an hour. Next, they were washed with 0.1% PBST (5 minutes, four times), and treated with 1X blocking solution for an hour. Then, the blocking solution was replaced with new blocking solution containing anti-digoxigenin-POD antibody (1:5000; Sigma-Aldrich, 11207733910) and 594-conjugated donkey anti-rabbit (IgG) antibody(1:1000; Jackson ImmunoResearch lab, 711-587-003), and the tissues were incubated for an hour at RT. After washing with PBS (5 minutes, four times), the tissues were incubated with Cy5-conjugated TSA (Perkin elmer, SAT705A001EA) for 5–20 minutes. The tissues were washed with 0.1% PBST (4 times, 5 minutes each) and mounted with Vectashield containing DAPI (Vector Laboratories, H-1200).

**Calcium imaging processing: registration and cell segmentation**

During the light sheet calcium imaging of GCaMP5g-P2A-PAmCherry transgenic mice, we observed tissue warping, such as shifting, swelling, and shrinking. Such movement was corrected by a custom non-rigid registration software written in Julia (*34*). Substantial differences in fluorescence intensity—for example, due to calcium influx—often caused abnormal registration. To reduce the impact of this issue, the imaging volumes before each stimulus delivery were chosen and median-filtered across 14 seconds (7 imaging volumes). These median-filtered imaging volumes were then registered to a fixed image, which was typically the median-filtered imaging volume in the middle time point of an imaging session. The resulting deformation vector was linearly interpolated over time to result in a deformation for each collected volume. After movement correction, we segmented cells responsive to any given ligand by a custom three-dimensional segmentation algorithm written in Julia. Because we used somatic calcium level as a measure of neuronal activity, the dendritic layer was intentionally removed from segmentation.

Ectopic-expression experiments were also performed by light sheet imaging. The VNE ectopically expressing *GCaMP-2A-VR* showed minimal baseline GCaMP fluorescence. Therefore, these imaging volumes were not subjected to registration. To segment responsive neurons, imaging volumes were max-projected over time and segmented by GCaMP intensity thresholding. Inspection suggests that segments contained 1–5 cells. To examine somatic GCaMP intensity, the dendritic layer was manually excluded from segmentation.

For each two-photon calcium imaging session, we acquired fewer than 200

$512 \times 512$ imaging frames and did not perform registration because we did not observe substantial tissue movement over this short time period. For the two-photon photoactivation accuracy test, calcium imaging data were segmented by the EM/MPM segmentation algorithm (*35*). When we attempted to identify genes expressed in the functional clusters, responsive neurons were segmented by a custom algorithm written in Julia (*36*). Overall, calcium imaging took less than 5 minutes and segmentation under 2 minutes.

#### Quantification and Statistical Analysis

##### Calcium imaging analysis: response and clustering

Calcium intensity traces were obtained from custom segmentation software written in Julia. To obtain neuronal response from the intensity trace, the time-weighted average was calculated from the 48 s of the post-stimulus period (*37*) and then the average of the 8 s pre-stim period was subtracted.

During light sheet calcium imaging, two separate aliquots of Ringer's solution were used as controls; one was used to select responsive cells and the other as a negative control. Neurons were discarded if their responses to at least one stimulus were not significantly higher than those evoked by one of the controls (4 trials, Student's *t*-test, $p$ value $< 0.01$) or if response intensities were below a manually-selected threshold intensity, which was approximately 10% higher than the average camera bias intensity.

For the reference functional clustering, responsive neurons from three VNEs were pooled and clustered by a variant of the mean shift clustering algorithm (*3*) (*38*). We initially obtained more than 80 clusters. We reasoned that our clustering algorithm detected subtle technical variations across experiments, categorizing what were biologically similar responses across animals into different clusters. These clusters were merged based on a pairwise penalty matrix. A penalty $P$ was calculated by the following equation:

$$P = c \times (1 + w \times f), \quad (1)$$

where $c$ indicates the Pearson correlation among clusters, $w$ is a weight, and $f$ indicates the Fano factor, which measures the variability of each cluster across different biological samples. $w$ was set to be 0.005. Pairs of clusters showing high $P$ were merged using manual supervision. In addition, one cluster contained two out of 5037 cells showing response to Ringer's control, and they were discarded from the analysis. Accordingly, we obtained 20 clusters, which are three more than the previously observed clusters (*3*). This difference might be attributed to several factors: In the present study, we used mice expressing GCaMP5g (and PAmCherry) while previous research used GCaMP3 mice, and more ligands and concentrations were used here.

During *post hoc* analysis of two-photon calcium imaging data (**Fig. S6A**), cells were clustered on the basis of the reference clustering: First, the mean response pattern of each reference cluster was calculated. Next, we measured pairwise cosine distance between each mean response and each cell's response obtained from two-photon calcium imaging. Each cell was assigned to the reference cluster showing the nearest response pattern to the cell. Due to small differences between the light sheet and two-photon experiments, the resulting clustering was further refined by repeating this process, using the cluster means that resulted from the initial two-photon clustering rather than the cluster means from the light sheet experiments.

##### Single cell analysis

Cel-seq2 analysis pipeline resulted in an UMI expression matrix (*30*). Briefly, sequencing reads were first demultiplexed by Cel-seq2 software. STAR aligner (ver 2.6.0a) aligned the sequencing reads to the mouse reference genome (GRCm38) using default parameters except for –sjdbOverhang, which was set to be 97 (*39*). A mouse genome annotation file (version M18, GTF file) was downloaded from Gencode. HTseq-count (ver 0.8.0) extracted the number of UMI for each gene using default parameters (*40*), with one exception for *Vmn1r237* described in Fig. S9. The resulting expression matrix was further analyzed in R using the Seurat single cell analysis package (*41*). Any gene with fewer than two UMIs was regarded as non-expressed. Cells were discarded from the analysis if they expressed fewer than 2000 or more than 8000 genes, or if more than 15% of UMIs corresponded to mitochondrial genes. For each cell, gene UMI counts were normalized by the total expression. The resulting values were multiplied by a scale factor 10,000 and log-transformed ($\log_e$).

To visualize read coverage, we used an R package, Gviz (*42*).

##### Phylogenetic analysis

A pairwise VR sequence distance matrix and the evolutionary relationship was obtained by Clustal Omega in the DNAStar software (*43*). The resulting phylogenetic tree was visualized using R (ggplot package (*44*)). The same distance matrix was used for classic multidimensional scaling analysis. This analysis was done in Julia (MultiVariateStats package).

#### Data and Software Availability

Software is available on GitHub, https://github.com/HolyLab. Single cell RNA sequencing results are on GEO (GSE127193). The extended *Vmn1r237* gene model used for read mapping in the present study is available in Data S1. The sequence of a *Vmn1r237* transcript variant is available in Genbank (MN473045).

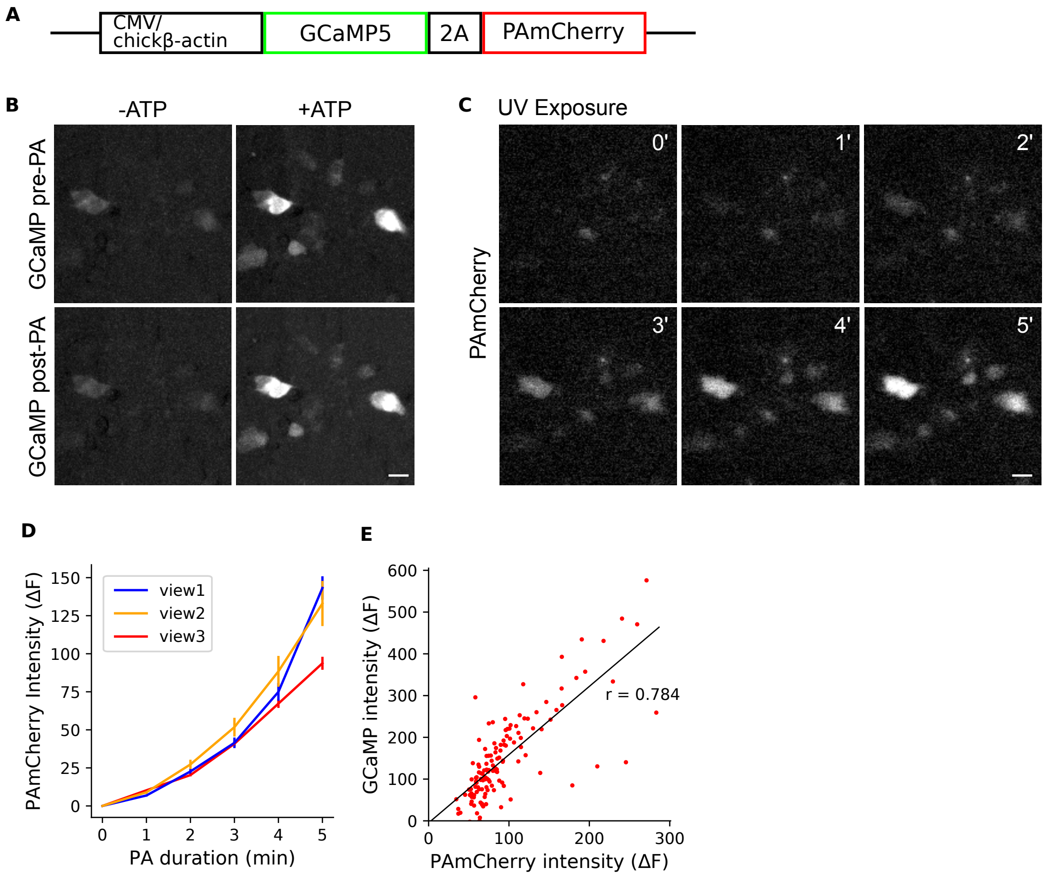


**Fig. S1. *In vitro* characterization of GCaMP-P2A-PAmCherry. (A)** The *GCaMP5g-2A-PAmCherry* DNA construct used for *in vitro* transfection. **(B)** GCaMP activity before (top) and after (bottom) photoactivation. To transiently increase intracellular calcium concentration, 10 $\mu$M ATP was delivered. Scale bar: 20 $\mu$m. **(C)** Representative images of photoactivated cells over time (405 nm, 1 mW). The same cells shown in (B) were visualized. **(D)** In three different fields of views (red, orange, blue), PAmCherry$^{high}$ cells were segmented after 5-minutes of photoactivation (0.3 J), and PAmCherry fluorescence intensity ($\Delta F$) was quantified as a function of photoactivation time. **(E)** After photoactivation, GCaMP was activated by adding 5 $\mu$M Ionomycin, a $Ca^{2+}$ ionophore, to the medium. From cells in the view 3 in (D, red), a quantification of activated GCaMP and PAmCherry signals ($\Delta F$) revealed a tight correlation (Pearson correlation coefficient $r$ = 0.784).

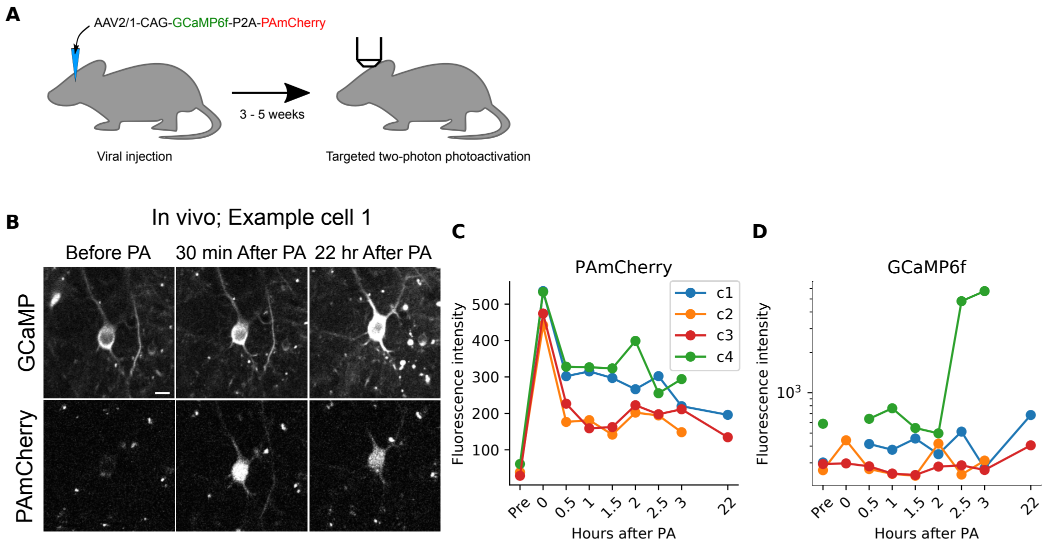


**Fig. S2.** ***In vivo* two-photon photoactivation.** **(A)** Injection of AAV2/1-CAG-GCaMP6f-P2A-PAmCherry and *in vivo* two-photon photoactivation. **(B)** The example shows GCaMP and PAmCherry signal at three different time points: before photoactivation, at 30 minutes, and at 22 hours after photoactivation. Scale bar: 10 $\mu$m. **(C and D)** From four cells individually photoactivated, **(C)** PAmCherry and **(D)** GCaMP fluorescence intensities were quantified across multiple time points.

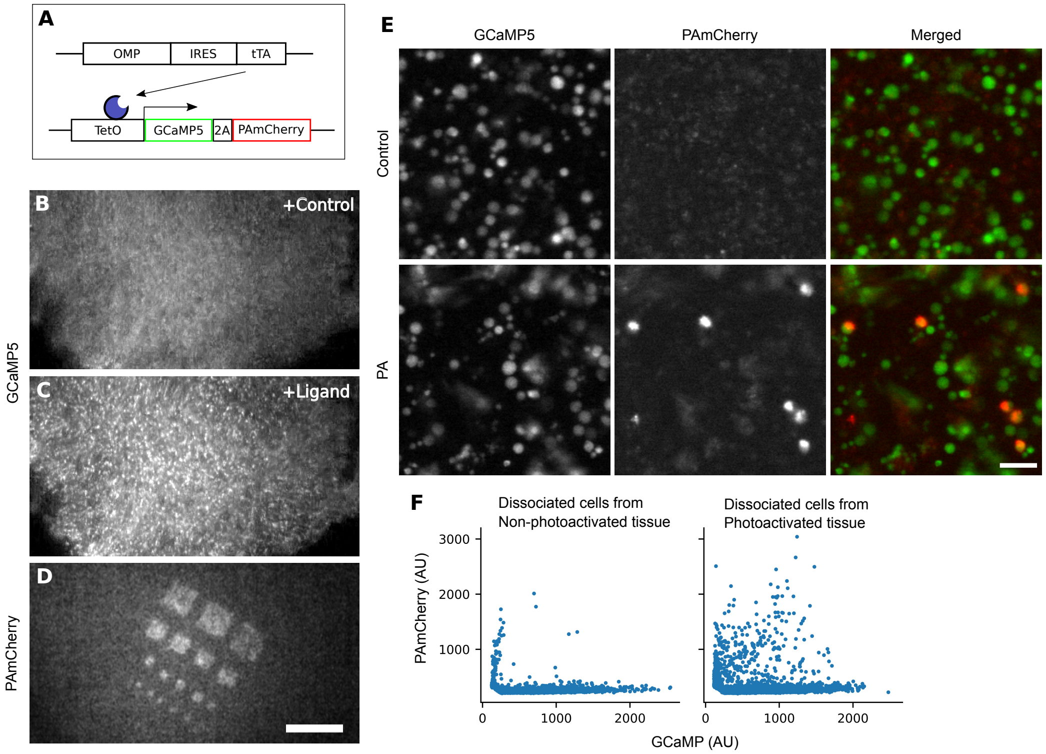


**Fig. S3. Macroscopic calcium imaging and photoactivation on an *ex vivo* VNE expressing GCaMP and PAmCherry (A)** Tet-off activator drives the expression of *GCaMP* and *PAmCherry* in mouse accessory olfactory sensory neurons. **(B and C)** Ligand application (10 $\mu$M E1050) increased GCaMP fluorescence. **(D)** Two-photon excitation at 810 nm increased PAmCherry signals in multiple square illumination regions ((100 $\mu$m)$^2$, (50 $\mu$m)$^2$, (20 $\mu$m)$^2$, and (13 $\mu$m)$^2$). Scale bar: 200 $\mu$m. **(E)** Representative images of dissociated VSNs from the non-photoactivated VNE (top row) or photoactivated VNE (bottom row). Scale bar: 40 $\mu$m. **(F)** Quantification of GCaMP and PAmCherry intensity of dissociated cells. Each dot indicates intensity of each cell.

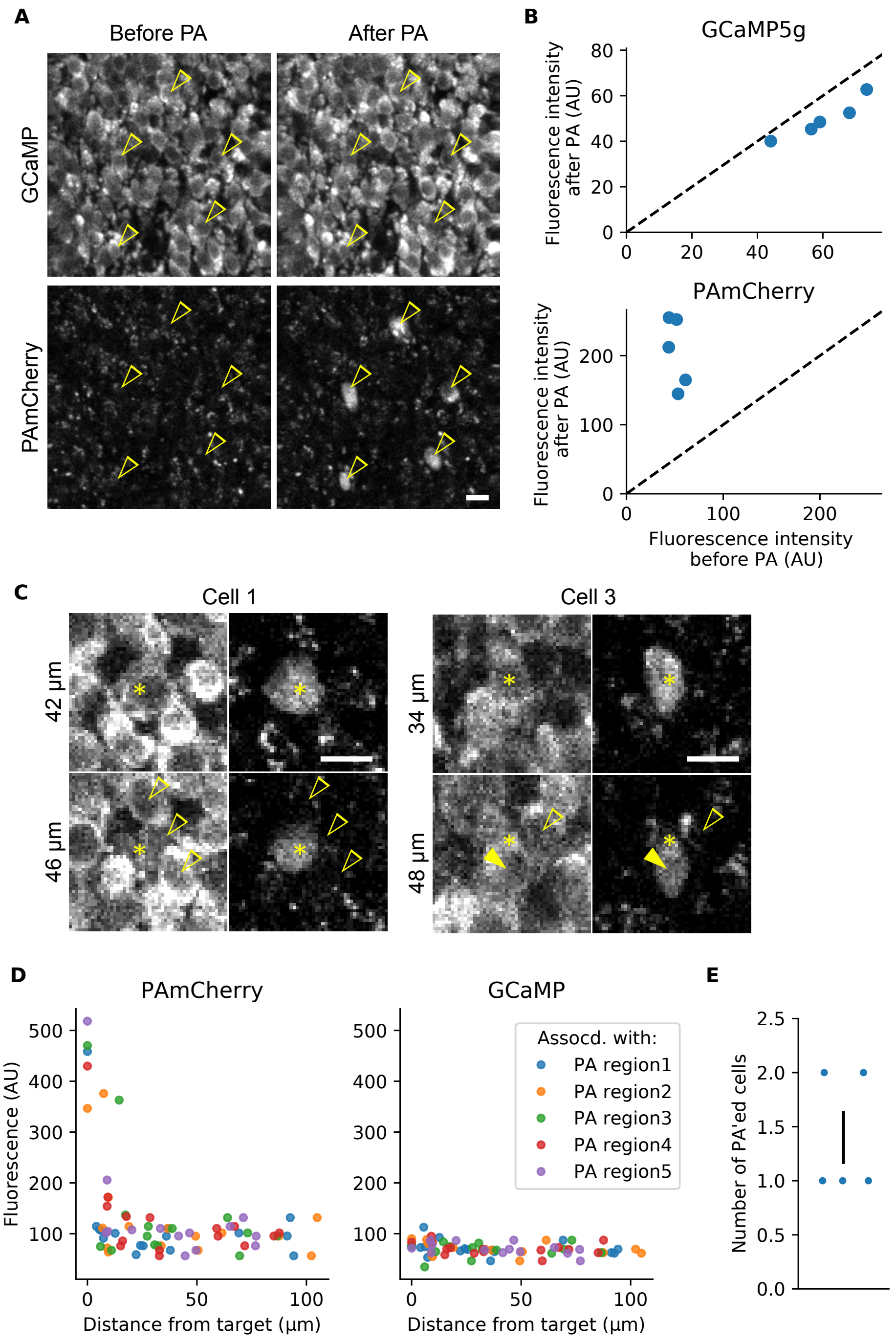
A
Before PA
After PA
GCaMP
PAmCherry
B
GCaMP5g
Fluorescence intensity after PA (AU)
PAmCherry
Fluorescence intensity before PA (AU)
C
Cell 1
Cell 3
42 µm
46 µm
34 µm
48 µm
D
PAmCherry
GCaMP
Fluorescence (AU)
Distance from target (µm)
Assocd. with:
PA region1
PA region2
PA region3
PA region4
PA region5
E
Number of PA'ed cells

**Fig. S4. Two-photon photoactivation provided near-single cell resolution photo-labeling in a dense population. (A and B)** GCaMP and PAmCherry intensity before and after photoactivation. **(A)** Representative images showing GCaMP and PAmCherry signals. Arrow heads indicate photoactivated neurons. Scale bar: 10 $\mu$m: **(B)** Quantification of GCaMP (top) and PAmCherry (bottom) fluorescence intensity before and after photoactivation. **(C–E)** Photoactivation accuracy was examined in 3-dimensional space. **(C)** Example images show two photoactivated target cells (cell 1 and cell 3). Vertical-axis labels indicate distance from the surface. Asterisks indicate photoactivation target cells; closed arrow heads are non-target cells with high PAmCherry signal. Open arrow heads indicate surrounding cells with background-level PAmCherry signal. Scale bar: 10 $\mu$m **(D)** PAmCherry (left) and GCaMP (right) intensity of example cells were measured as a function of the euclidean distances from PA target cells. Each dot indicates one cell, where those of the same color are neighbors of the same target PA region. Cells at 0 $\mu$m are the target cells. The green and blue dots at 0 $\mu$m were jittered along the vertical axis to avoid overlap. **(E)** The number of PAmCherry$^{high}$ cells associated with their PA region shown in panel (A). The error bar indicates s.e.m.

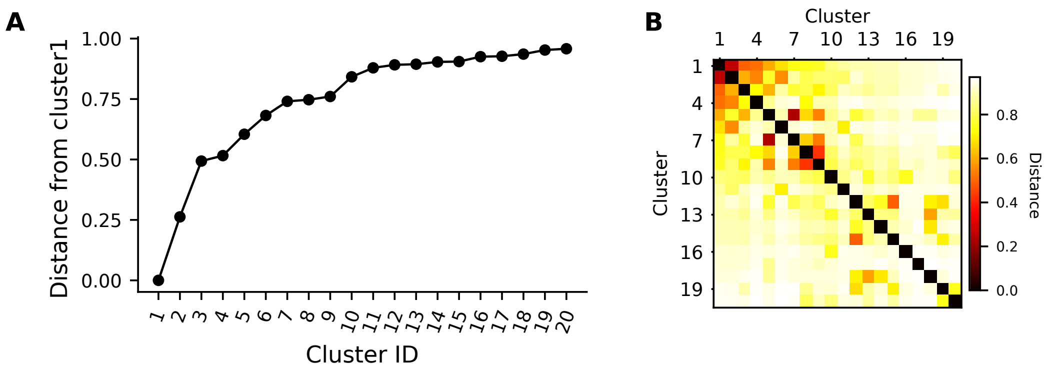


**Fig. S5. The similarity among functionally-defined VSN types.** Normalized euclidean distance **(A)** between cluster1 and the others and **(B)** among all clusters shown in the reference clustering (**Fig. 2B**). The colorbar indicates normalized euclidean distance.

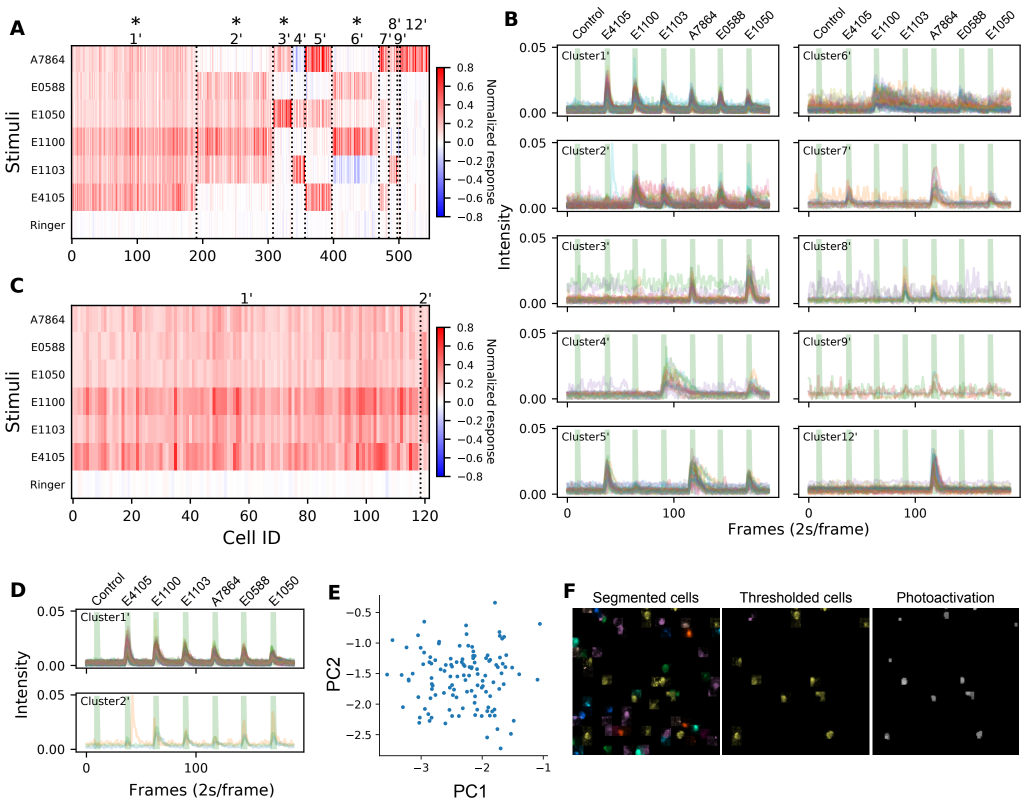


**Fig. S6.** ***Post hoc*** **analysis of two-photon calcium imaging. (A)** The raster plot shows VSN response to six ligands (A7864, E0588, E1050, E1100, E1103, and E4105; and Ringer control). The concentration of each ligand was 10 $\mu$M. The colorbar indicates normalized response. Based on the reference clustering, neurons were clustered into 10 clusters (see Methods). Cluster1$'$ cells responded to all six ligands, showing the same response behavior as cluster1 in the reference clustering; moreover, this cell type was the most abundant among responsive neurons, again consistent with the reference. The response patterns of other cell types, cluster2$'$–cluster9$'$, and cluster12$'$, also matched the reference types. Accordingly, these results suggested that two-photon calcium imaging with a selected stimulus set allows for the identification of cell types of interest. **(B)** Response traces of all neurons shown in (A). Each colored line indicates a response trace from one cell. GCaMP intensity was measured by $\Delta F$. Green bars show timing of ligand delivery. Control solution, E4105, E1100, E1103, A7864, E0588, and E1050 were applied for 5 seconds in order in this particular example. **(C–F)** An example of cell selection by thresholding. Cluster1$'$ cells were chosen as those exceeding a threshold to every ligand. Their cluster identities and response profile were illustrated by **(C)** a raster plot or **(D)** a temporal trace plot. We obtained $\sim$120 cells, among which more than 95% were from cluster$1'$ on the basis of a more laborious *post hoc* analysis. **(E)** A PCA analysis of temporal traces of the chosen cells revealed only a single cluster, indicating that their response patterns are homogeneous. **(F)** Example images of cell segmentation (left), cell selection (middle), and photoactivation mask (right). Cells responsive to any ligand were automatically segmented (left), where different colors indicate different functional types. Among segmented cells, those responsive to all ligands were selected by thresholding (middle), and a photoactivation mask was created (right).

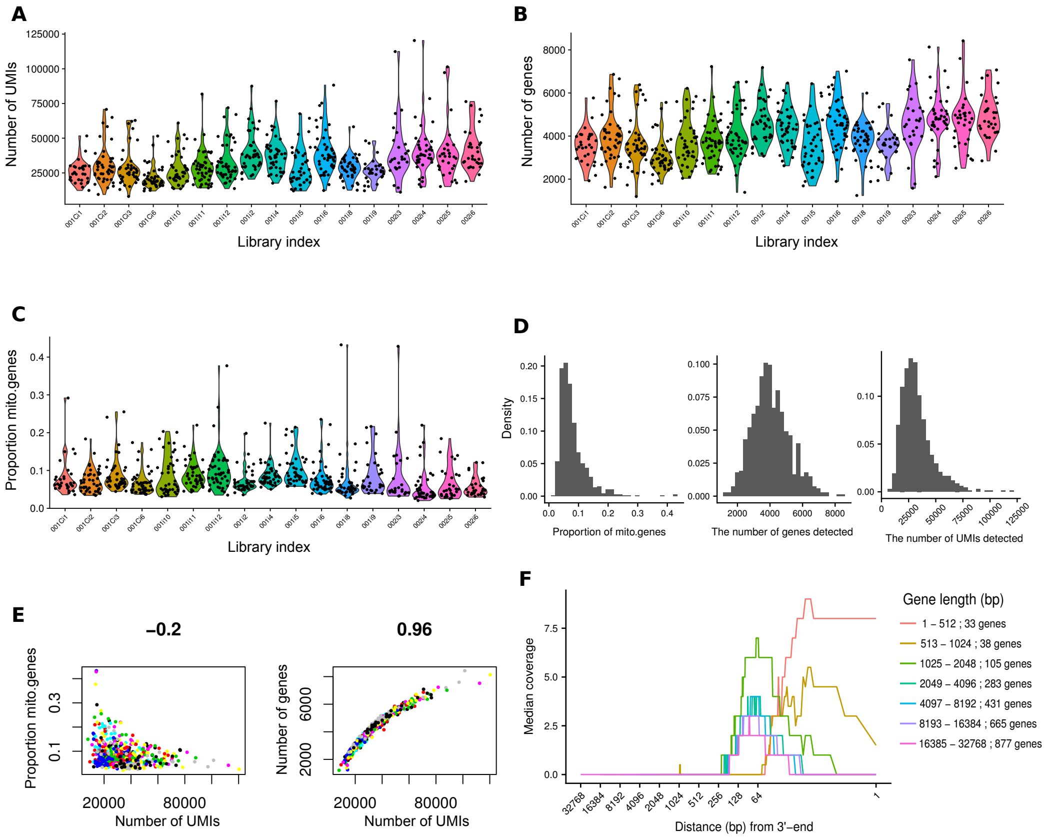


**Fig. S7. scRNAseq quality control.** **(A–C)** Violin plots showing **(A)** UMI counts in each cell, **(B)** the number of genes in each cell, and **(C)** the proportion of mitochondrial gene expression in each cell. The horizontal axis shows Illumina sequencing library indices, which tag different sequencing library preparations. **(D)** Histograms of total population for quantities plotted in (A–C). **(E)** Scatter plots show a slight negative correlation (-0.2) between UMI counts and the proportion of mitochondrial genes (left), and strong positive correlation (0.96) between UMI counts and the number of genes detected (right). **(F)** Coverage plot with the median read coverage as a function of distance from the 3' end. Only uniquely aligned reads were counted. Different line colors indicate different groups of genes; genes were grouped by their length.

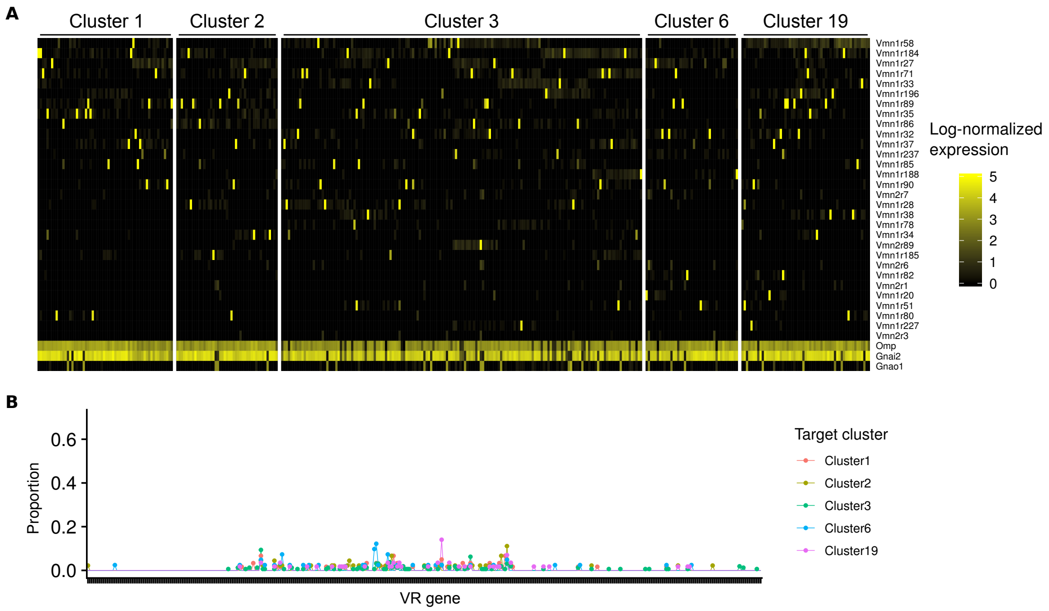


**Fig. S8. PhOTseq control cells were not enriched with a specific VR type. (A)** Together with PA'ed cells shown in Fig. 2C, non-photoactivated cells were sequenced. The list of genes exhibited are the same as that shown in Fig. 2C: The top 30 highly expressed VR genes when averaged across all cells sequenced, markers for VSN (*Omp*), V1R family (*Gnai2*), and V2R family (*Gnao1*). **(B)** The proportion of a VSN type in each experimental control group shown in (A). Each tick along the horizontal axis represents a different VR gene. The vertical axis shows the proportion of cells for which this gene was the highest-expressed VR gene. Line color indicates the target cluster.

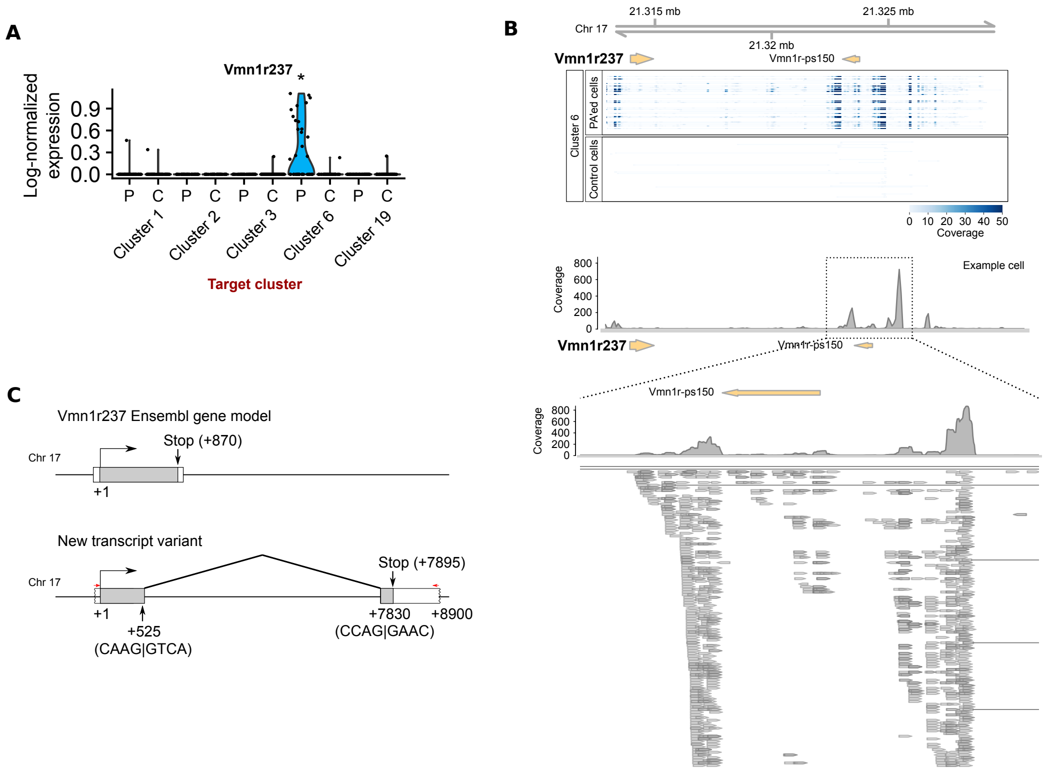


**Fig. S9.** ***Vmn1r237*** **transcript variants.** **(A)** The violin plot shows the normalized number of reads aligned to the original *Vmn1r237* Ensemble gene model. Using this gene model, *Vmn1r237* was $\sim$50$\times$ less abundant in terms of mRNA copy number than VR genes identified from other clusters (see Fig. 2E). **(B)** Read coverage on the *Vmn1r237* locus. The *Vmn1r237* gene model was mapped onto mouse chromosome 17 (top). *Vmn1r-ps150* was located roughly 10 kb downstream of *Vmn1r237*. *Vmn1r237* and *Vmn1r-ps150* are encoded in the sense and the anti-sense strand, respectively (arrow direction). The read coverage heatmap was obtained from cluster6-targeted cells (middle top; 39 photoactivated and 41 non-photoactivated control cells). In the majority of PA'ed cells, most reads were aligned to $\sim$10kb downstream of *Vmn1r237*. In the bottom half of (B), the read coverage map of an example PA'ed cell is shown. The enlarged view shows the direction of reads aligned in this region. The direction of the majority of reads was the same as that of *Vmn1r237*, possibly suggesting that these reads are originated from an unidentified 3' UTR of *Vmn1r237*. **(C)** A transcript variant of *Vmn1r237* was cloned and sequenced, providing an evidence of a 10kb-long 3'-

UTR of *Vmn1r237*. Red arrows show PCR primers used for cloning. Gray boxes indicate exons, white boxes indicate introns. The boundaries of introns are wavy if they are likely to be further extended. The black arrows and +1 indicate the translation start site. The nucleotide sequences show potential splicing junctions. When these reads were included, *Vmn1r237* was expressed at levels similar to that of other VR genes shown in Fig. 2E.

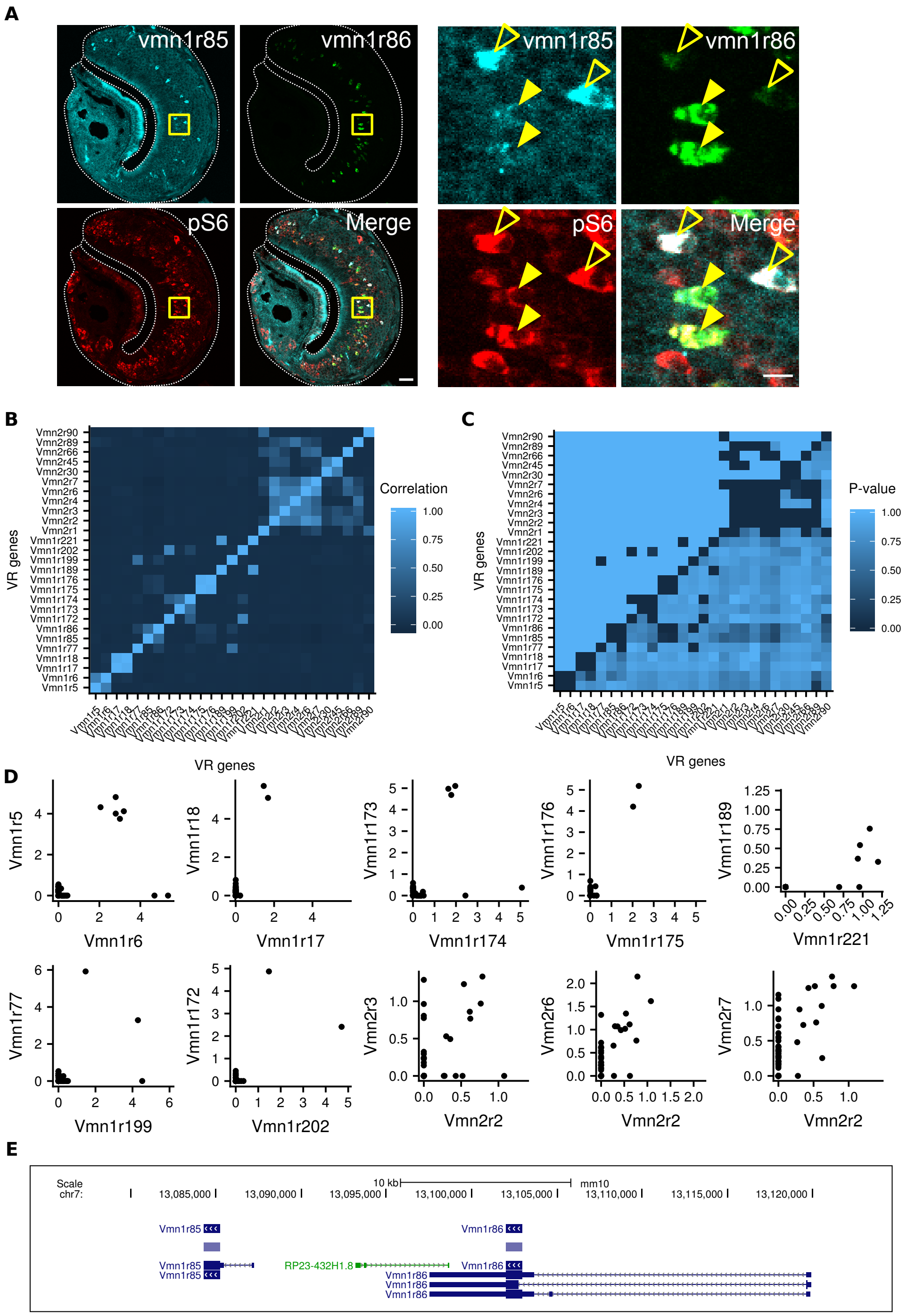

A
vmn1r85
vmn1r86
pS6
Merge
B
Correlation
VR genes
C
P-value
D
VR genes
E
Scale
chr7:
10 kb
mm10
Vmn1r85
Vmn1r86
RP23-432H1.8

**Fig. S10. Expression of multiple VR genes in single cells. (A)** Representative images of double fluorescent in situ hybridization (FISH) and pS6 immunohistochemistry (IHC) on a VNO section (left) and their zoomed-in views (yellow boxes; right). Riboprobes for *Vmn1r85* (cyan) and *Vmn1r86* (green) were applied during FISH. pS6 indicates recently activated neurons: Before FISH and IHC, the animal was treated with 50 $\mu$M E1050 *in vivo*. Forty minutes after the ligand treatment, the VNO was dissected out and subjected to double FISH and IHC. In the zoomed-in views (right), filled arrow heads indicate cells showing high *Vmn1r86* but low *Vmn1r85* signals. Open arrow heads indicate cells showing high *Vmn1r85* but low *Vmn1r86*. Note that we cannot exclude the possibility that cross-hybridization and/or fluorescence cross-talk (bleed-through) contributes to this result. **(B)** Heatmap showing pairwise correlations of VR gene expression across cells. Only those VR genes expressed abundantly in more than 3 cells were included, and VR pairs showing Pearson correlation coefficient $r > 0.45$ are displayed. **(C)** Heatmap of $p$ values associated with the pairwise correlations in (B). Raw $p$ values are shown below the diagonal. Adjusted $p$ values by Holm's method are reported above the diagonal. **(D)** Among the correlated VR genes shown in (B), log-normalized expression from all V1R (*Vmn1r*) with $r > 0.45$ and three example V2R (*Vmn2r*) pairs are plotted. In the top row, VR gene pairs in the first four plots show consistent high/low coexpression, and they are adjacent to each other in the genome; the horizontal axis indicates the expression of downstream genes. As with *Vmn1r86* and *Vmn1r85*, some pairs (*Vmn1r5*/*Vmn1r6* and *Vmn1r173*/*Vmn1r174*) exhibited selective expression of the downstream gene in particular cells. The second row shows gene pairs that were reported as correlated but which do not show consistent clusters of high/low coexpression. **(E)** Ensemble gene models show the chromosomal location of *Vmn1r86* and *Vmn1r85*.

**Movie S1. Example calcium response and online cell segmentation.** Example calcium response to conrol, E1100, E1103, A7864, E0588, and E1050 (left). Concentration of each ligand was 10 $\mu$M. Responsive neurons were segmented online and randomly pseudocolored (right). Images were acquired at 0.5 Hz. Video plays at 20 frames per second.
**Movie S2. Stimulus responses under ectopic expression of *Vmn1r237*.** Single plane extracted from light sheet imaging volume of wildtype mice infected with rAAV2/8-CAG-GCaMP6f-2A-Vmn1r237. 60 planes were collected at a rate of one volume every two seconds. Video plays at 20 frames per second. Stimulus labels in top left at their time of application. The apical layer is the sensory surface of the tissue.

### References


1. H. Markram, *et al.*, *Nature Reviews Neuroscience* **5**, 793 (2004).
2. M. Nassar, *et al.*, *Frontiers in Neural Circuits* **9**, 1 (2015).
3. P. S. Xu, D. Lee, T. E. Holy, *Neuron* **91**, 878 (2016).
4. H. Okuno, *Neuroscience Research* **69**, 175 (2011).
5. S. Qiu, *et al.*, *Frontiers in Genetics* **3**, 1 (2012).
6. S. Haga-Yamanaka, *et al.*, *eLife* **2014**, 1 (2014).
7. J. Fuzik, *et al.*, *Nature Biotechnology* **34**, 175 (2015).
8. C. R. Cadwell, *et al.*, *Nature Biotechnology* **34**, 199 (2015).
9. B. F. Fosque, *et al.*, *Science* **347**, 755 (2015).
10. J. Akerboom, *et al.*, *The Journal of Neuroscience* **32**, 13819 (2012).
11. T.-W. Chen, *et al.*, *Nature* **499**, 295 (2013).
12. F. V. Subach, *et al.*, *Nature Methods* **6**, 153 (2009).
13. M. D. Ryan, A. M. King, G. P. Thomas, *Journal of General Virology* **72**, 2727 (1991).

14. K. Wilson, G. Raisman, *Brain research* **185**, 103 (1980).

15. D. Keller, C. Erö, H. Markram, *Frontiers in neuroanatomy* **12** (2018).

16. T. Bozza, P. Feinstein, C. Zheng, P. Mombaerts, *Journal of Neuroscience* **22**, 3033 (2002).

17. P. Feinstein, T. Bozza, I. Rodriguez, A. Vassalli, P. Mombaerts, *Cell* **117**, 833 (2004).

18. E. A. Hallem, M. G. Ho, J. R. Carlson, *Cell* **117**, 965 (2004).

19. X. Grosmaitre, A. Vassalli, P. Mombaerts, G. M. Shepherd, M. Ma, *Proceedings of the National Academy of Sciences* **103**, 1970 (2006).

20. P. Mombaerts, *Nature Reviews Neuroscience* **5**, 263 (2004).

21. J. Loconto, *et al.*, *Cell* **112**, 607 (2003).

22. B. Stein, M. T. Alonso, F. Zufall, T. Leinders-Zufall, P. Chamero, *PLoS ONE* **11**, 1 (2016).

23. Y. Isogai, *et al.*, *Nature* **478**, 241 (2011).

24. B. von der Weid, *et al.*, *Nature Neuroscience* **18**, 1455 (2015).

25. Y. Jiang, *et al.*, *Nature Neuroscience* **18**, 1446 (2015).

26. P. S. Xu, T. E. Holy, *Pheromone Signaling* (Springer, 2013), pp. 201–210.

27. T. F. Holekamp, D. Turaga, T. E. Holy, *Neuron* **57**, 661 (2008).

28. A. Kaur, S. Dey, L. Stowers, *Pheromone Signaling* (Springer, 2013), pp. 189–200.

29. H. A. Arnson, X. Fu, T. E. Holy, *Journal of visualized experiments: JoVE* **37** (2010).

30. T. Hashimshony, *et al.*, *Genome Biology* **17**, 1 (2016).

31. S. E. G. Lampe, B. K. Kaspar, K. D. Foust, *Journal of visualized experiments: JoVE* **93** (2014).

32. K. Kang, D. Lee, S. Hong, S.-G. Park, M.-R. Song, *Journal of Biological Chemistry* **288**, 2580 (2013).

33. P. A. Gray, *et al.*, *Journal of Neuroscience* **30**, 14883 (2010).

34. T. E. Holy, Blockregistration package, `https://github.com/HolyLab/BlockRegistration` (2019).

35. M. L. Comer, E. J. Delp, *IEEE Transactions on image processing* **9**, 1731 (2000).

36. T. E. Holy, Cellsegmentation package, `https://github.com/HolyLab/CellSegmentation` (2019).

37. D. Turaga, T. E. Holy, *The Journal of Neuroscience* **32**, 1612 (2012).

38. T. E. Holy, Neighborhoodclustering package, `https://github.com/HolyLab/NeighborhoodClustering.jl` (2019).

39. A. Dobin, *et al.*, *Bioinformatics* **29**, 15 (2013).

40. S. Anders, P. T. Pyl, W. Huber, *Bioinformatics* **31**, 166 (2015).

41. A. Butler, P. Hoffman, P. Smibert, E. Papalexi, R. Satija, *Nature biotechnology* **36**, 411 (2018).

42. F. Hahne, R. Ivanek, *Statistical Genomics* (Springer, 2016), pp. 335–351.

43. F. Sievers, *et al.*, *Molecular systems biology* **7**, 539 (2011).

44. G. Yu, D. K. Smith, H. Zhu, Y. Guan, T. T.-Y. Lam, *Methods in Ecology and Evolution* **8**, 28 (2017).
